# A modular intranasal *Nm*-nOMV RBD concatemer vaccine induces broad betacoronavirus-neutralizing antibodies

**DOI:** 10.64898/2026.11.26.769503

**Authors:** Kaneemozhe Harichandran, Wouter J.P. van den Braak, Ruben J.G. Hulswit, Joline van der Lee, Niklas Krug, Franziska Retschlag, Dinja Oosterhoff, Christian Drosten, Marcel A. Müller, Berend-Jan Bosch, Corine P. Kruiswijk

## Abstract

Betacoronaviruses pose a major threat to human and animal health due to their propensity to cause spillover infections. Broadly protective vaccines that induce mucosal immunity may strengthen pandemic preparedness by limiting infection and onward transmission. Here we have developed intranasal betacoronavirus vaccine candidates combining AdJane Biosciences’ *Neisseria meningitidis* native outer membrane vesicle (*Nm*-nOMV) platform, a self-adjuvating mucosal delivery platform recently shown to be safe and well tolerated in a first-in-human Phase 1 trial, with multivalent coronavirus spike immunogens. We evaluated betacoronaviruses full Spike proteins, chimeric full Spike proteins, and Receptor Binding Domain (RBD) concatemers representing sarbecoviruses, merbecoviruses, and embecoviruses. Antigenic profiling identified immunogens retaining activity against human convalescent sera and characterized monoclonal antibodies. In mice, intranasal immunization with *Nm*-nOMVs combined with RBD concatemers induced mucosal IgA responses, systemic IgG and broad neutralization of homologous and heterologous human and bat related betacoronaviruses. Together, findings from the study identify multivalent RBD concatemers co-administered with the *Nm*-nOMV platform as a promising, modular antigen design for the development of broadly protective mucosal vaccines against emerging betacoronaviruses.

## Introduction

Coronaviruses (CoVs) are enveloped positive-sense RNA viruses occurring globally in birds and mammals^1^. CoVs have a tendency for cross-species transmission and subsequent emergence in immunologically naïve host populations^1,2^. This risk is further amplified by increasing globalization and human encroachment into wildlife habitats, highlighting the persistent threat of emerging CoVs to both human and animal health and the need for effective pandemic preparedness measures. Since 2002, three major zoonotic coronaviruses—SARS-CoV, MERS-CoV, and SARS-CoV-2—all belonging to the genus Betacoronavirus (BCoV), have emerged and caused significant human outbreaks, highlighting the recurrent threat of zoonotic coronavirus spillover^3,4^. Consequently, for sustainable pandemic preparedness, future vaccine strategies should provide protection beyond currently circulating CoVs.

The coronavirus spike (S) protein is the primary target for vaccine development^5^. This class I fusion glycoprotein forms a homotrimer comprising an ectodomain, transmembrane domain, and intracellular tail^6^. The ectodomain is divided into S1, which contains the receptor-binding domain (RBD), and S2, which mediates membrane fusion^6^. In SARS-CoV-2, the RBD binds angiotensin-converting enzyme 2 (ACE2) and is a major target of serum neutralizing antibodies, making it an attractive immunogen for inducing neutralizing and cross-neutralizing responses^7^. The RBD region transitions between a closed (‘down’) and receptor-accessible (‘up’) conformation, modulating antibody accessibility to specific epitopes^8,9^. Despite extensive antigenic evolution of the RBD during the pandemic and the emergence of variants capable of escaping pre-existing immunity^10,11^, cross-protective antibodies targeting conserved RBD epitopes can neutralize multiple SARS-CoV-2 variants and related betacoronaviruses, including SARS-CoV and bat SARS-CoV WIV-1^12,13^

Several studies have demonstrated that systemic vaccination with combinations of coronavirus S proteins or RBDs can induce broadly reactive and cross-neutralizing antibodies in mice^7,14,15^. Intramuscular vaccination leads to the development of serum and tissue-resident IgG antibodies combating the virus after infection and preventing serious pathology and disease. However, these intramuscular vaccines do not fully protect from (re)infection and can reduce, but not prevent, onward transmission^15,16^. Human CoVs (HCoV) initiate and/or efficiently replicate in the upper respiratory tract (URT). In contrast to intramuscular vaccinations, intranasal vaccines can induce local innate as well as adaptive mucosal immune responses in the URT by eliciting high neutralizing antibody titers (especially IgA) at the site of viral replication^17,18^. IgA can cross the luminal side of the mucosal epithelial lining, where they are able to neutralize pathogens^19,20^.

Previous studies have demonstrated that intranasal CoV vaccination can elicit robust mucosal IgA responses and confer protection against infection^21–23^. A promising platform for intranasal vaccine development is based on *Neisseria meningitidis (Nm)* derived native Outer Membrane Vesicles (nOMVs), non-infectious vesicles naturally released by Gram-negative bacteria that possess intrinsic immunostimulatory properties through pathogen-associated molecular patterns (PAMPs), including lipopolysaccharides and lipoproteins. *Nm*-nOMV-based vaccines have demonstrated efficacy in preclinical studies against several viral antigens, including respiratory syncytial virus and hepatitis B surface antigen^24,25^. Importantly, we recently demonstrated the safety and tolerability of an *Nm*-nOMV-based SARS-CoV-2 vaccine in a first-in-human Phase 1 clinical trial providing support for the use of this platform for intranasal vaccine development^26^.

In this manuscript we have combined antigenic breadth with mucosal delivery by developing an intranasal *Nm*-nOMV platform incorporating diverse betacoronavirus S immunogens. We compared full S ectodomains, chimeric S proteins, and multivalent RBD concatemers representing sarbecoviruses, merbecoviruses, and embecoviruses (ECoV) for immunogenicity and cross-neutralizing activity in mice. Following antigenic characterization, selected immunogens were evaluated for immunogenicity and cross-neutralizing activity after intranasal vaccination in mice. This approach enabled us to identify multivalent RBD concatemers as a compact and modular antigen format capable of inducing broad neutralizing antibody responses against antigenically diverse betacoronaviruses.

## Results

### Vaccine antigen design

With the aim of generating a vaccine that elicits a broad antibody responses to multiple *Betacoronavirus*es, we selected S proteins from eleven members of human-infecting *Betacoronavirus*es and related bat-borne counterparts spanning the *Embeco-*(HCoV-OC43 A and B, HCoV-HKU1 A and B), *Merbeco-*(MERS-CoV, HKU4, PDF-2180), and *Sarbecovirus* (SARS-CoV, WIV-1, SARS-CoV-2, RaTG13) subgenera (**Figures 1A and 1B**). Three types of antigens were included in the designs: intact S protein ectodomains, intra-subgenera chimeras thereof, in which RBD and the residual S protein components were combined, and individual or concatenated RBDs (**Figure 1C**). The designed constructs were synthesized, used for transient expression in CHO Express cells, and purified by IMAC and SEC. Of the initial collection of 65 antigens, 43 antigens were expressed at sufficiently high levels (>0.15 mg/mL) for subsequent analyses.

**Figure 1:**
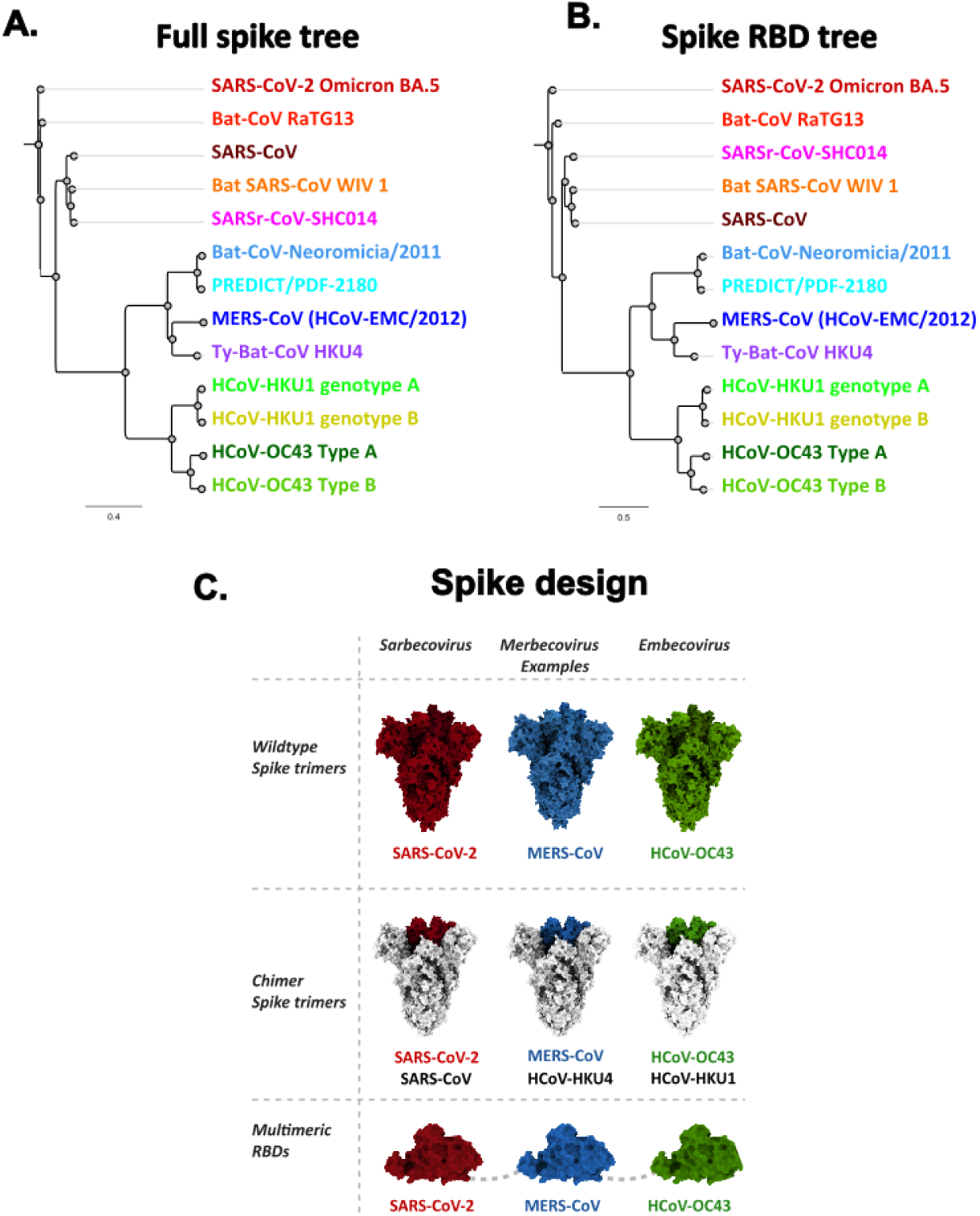
***Betacoronavirus phylogeny and vaccine candidate design.* A**) Phylogenetic tree of betacoronavirus full spike proteins used in designing of vaccine candidates. Sarbeco virus spikes are represented in shades of red, merbeco virus spikes in shades of blue and embeco virus spikes in green for easier perception of the data. **B**) Phylogenetic tree of betacoronavirus spike protein RBDs used in designing of vaccine candidates. **C**) Illustration showing examples of designed betacoronavirus full spikes, chimeric spike and RBD concatemers.

### Antigenic profiling of vaccine candidates using human coronavirus-reactive sera

To be able to consider the human background immunity in the vaccine antigen design, we first characterized antibody responses in paired serum samples collected from 20 individuals before (V1) and after (V3) SARS-CoV-2 vaccination (**Figure 2A**). Antibody endpoint titers against the four endemic HCoVs, SARS-CoV-2, and MERS-CoV were determined using an immunofluorescence assay (IFA)^27^ (**Supplement Figure 1A-F**). Based on the antibody reactivity results, seven sera exhibiting the highest antibody titers against all representative CoVs were selected and used to screen against 43 synthesized proteins by indirect ELISA with serum dilutions from 1:200–1:6,400 (**Figure 2B, Supplement Table 1**). In addition to the above-mentioned sera, a control serum with high titers for SARS-CoV-2 that was taken from an infected and vaccinated patient was included to monitor the effect of hybrid immunity upon SARS-CoV-2 vaccination and infection. Furthermore, we also included a SARS-CoV antibody reactive serum that was collected three and a half years after a SARS-CoV Frankfurt strain infection. Results showed that all sera collected before SARS-CoV-2 vaccination, V1, recognized only *Embecovirus* full and chimeric S protein trimers (**Figure 2B, Supplement Table 1**). Sera taken post-SARS-CoV-2 vaccinations, V3, reacted with full and chimeric S protein trimers and multimeric RBD proteins representing *Sarbeco-, Embeco-*, and most *Merbecoviruses*. Furthermore, V3 sera also showed an increase in antibody recognition towards endemic CoV S proteins, indicating cross-reactivity (**Figure 2B**). In addition, antibodies elicited after SARS-CoV-2 vaccination also cross-reacted with MERS-CoV and HKU4 S proteins (**Figure 2B**). From the cross-reactivity profiles and protein expression levels (**Supplement Table 2**), three full S, three chimeric S, and six RBD concatemer proteins were selected as candidates for immunogenicity studies.

**Figure 2:**
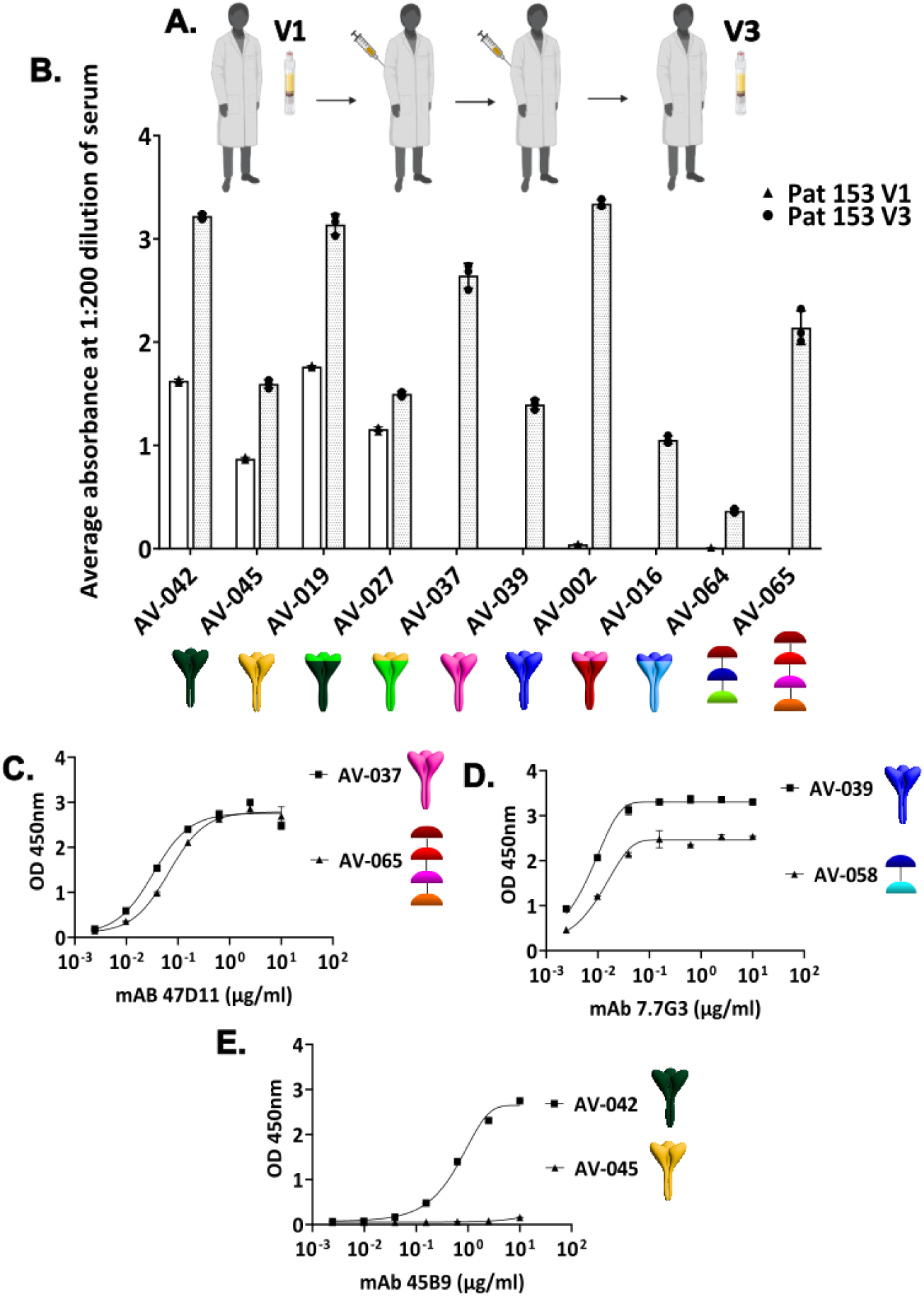
Antigenic profiling using human coronavirus-reactive sera and antigenic validation using coronavirus spike-specific monoclonal antibodies. A) Illustration showing the timeline of sera collection from healthcare workers. V1-before SARS-CoV-2 vaccination, V3-after 2 doses of SARS-CoV-2 vaccination. **B**) Graph showing a subset (10 out of 43) of results from vaccine candidate screening ELISA using defined human-coronavirus-reactive serum Pat 153 V1 and V3. The x-axis represents a subset of full, chimeric spikes and RBD concatemers that were screened, and the y-axis represents OD 450nm absorbance at 1:200 dilution of the sera. The V1 time point is represented by clear bars, and the V3 time point is represented by shaded bars. **C**) Graph showing vaccine candidate screening ELISA using mAb 47D11 (conformational Ab against SARS-CoV-2 S1B/RBD region). The x-axis represents dilution of antibodies, and the y-axis represents OD value at 450 nm. **D**) Graph showing vaccine candidate screening ELISA using mAb 7.7G3 (conformational Ab against MERS-CoV S1B/RBD region). The x-axis represents dilution of antibodies, and the y-axis represents OD value at 450 nm. **E**) Graph showing vaccine candidate screening ELISA using mAb 45B9 (conformational Ab against HCoV-OC43 strains, BCoV, and ECoV S1B/RBD region). The x-axis represents dilution of antibodies, and the y-axis represents OD value at 450 nm.

### Antigenic validation using coronavirus spike-specific monoclonal antibodies

To assess whether the designed His-tagged immunogens displayed a native folding, we tested the purified recombinant proteins against a panel of previously published^28–30^ monoclonal antibodies specific to different *Betacoronavirus* S proteins (**Supplement Table 3**). Immunogens were coated on the ELISA plates and probed against the panel of mAbs. An anti-His mAb was used as a positive control for all immunogens. In all cases, the immunogens were recognized in a concentration range of 0.002–10 µg/ml by the respective S-protein-specific monoclonal antibodies (mAb) except for AV-045, a full S protein of HCoV-HKU1 genotype B^28^ (**Figure 2C-E, Supplement Figure 3A-C**). These results support preservation of antigenically relevant epitopes in the recombinant immunogens and of the designed immunogens.

### First mice immunogenicity study

Selected vaccine candidates (**Table 1**) were used to twice immunize (Day 0 and Day 21, **Figure 3A**) mice (n=10 per group) and were sacrificed on Day 35. All applied vaccine candidates with their corresponding naming are listed in **Table 1**. Sera from mice were pooled and subjected to neutralization tests (NT) against four Vesicular Stomatitis virus pseudoparticles (VSVpp)-carrying S proteins of SARS-CoV-2 BA.5 (S-CoV-2, red), SARS-CoV (S-CoV, brown), bat-borne SARS-related CoV WIV1 (Sr-CoV, orange), and MERS-CoV (M-CoV, blue). As a control, full S proteins of SARS-CoV-2 BA.5, SARSr-CoV SHC014 and MERS-COV were used to intranasally vaccinate mice (**Figure 3A**). S-CoV VSVpp was exclusively used as a cross-reactivity control, as the corresponding S protein was not included as a vaccine antigen. Results from these neutralization assays showed expected cross-neutralization patterns (**Figure 3B-C**). S-CoV-2 S protein-vaccinated pooled mice sera showed no cross-neutralization with S-CoV, which emerged in 2003. However, mice vaccinated with the S-CoV S-related protein SARSr-COV SHC014 showed cross-neutralization against all *Sarbeco* VSVpp with the highest titers for closely related Sr-CoV (NT50=1,280; **Figure 3C**) (**Figure 3B–C**). No cross-neutralization was observed between *Sarbeco* and M-CoV S-VSVpp and vice versa. Calculated NT50 values for the NT assays are represented in **Figure 3C**.

**Figure 3:**
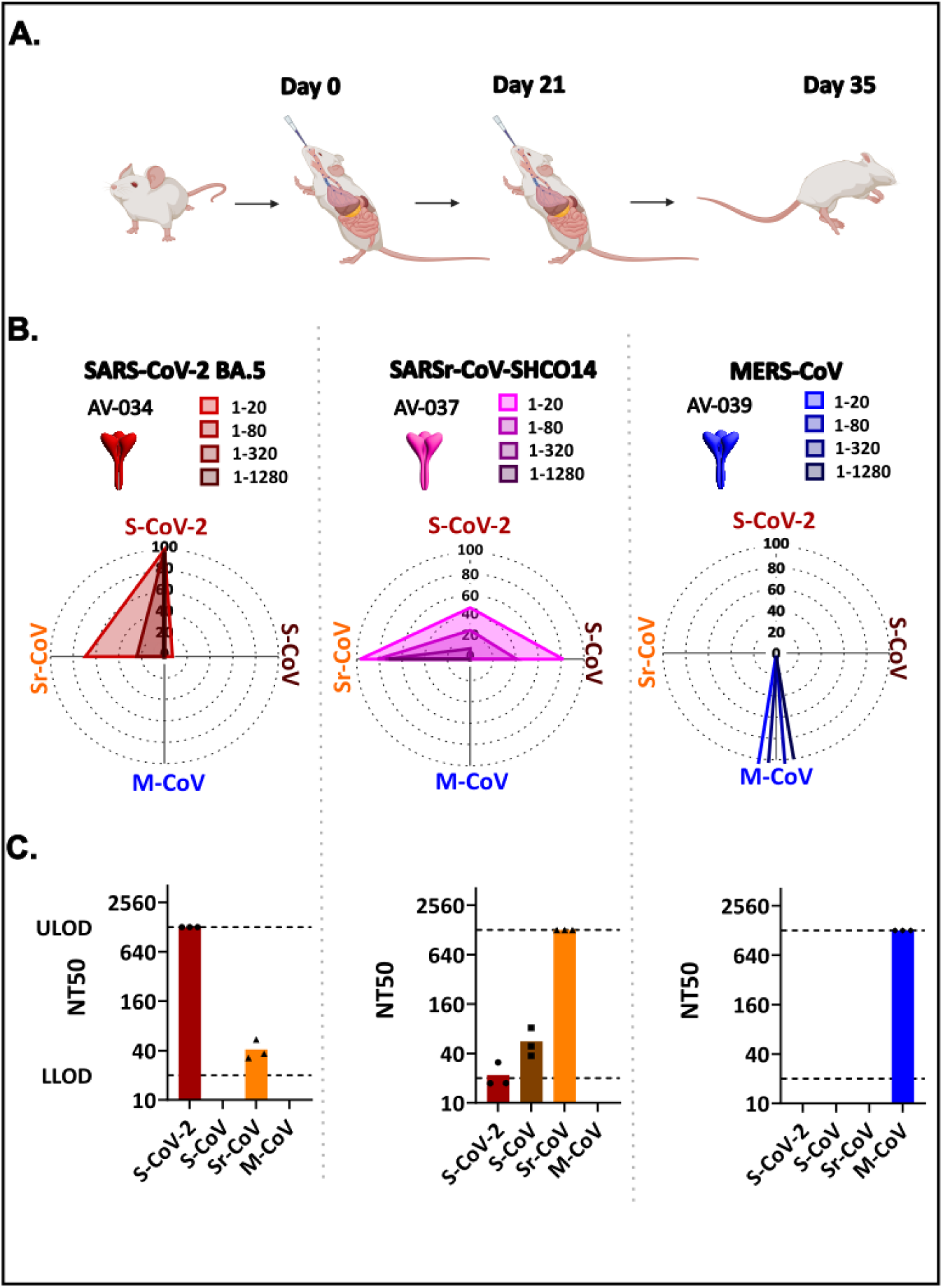
***First mice immunogenicity study -full spike groups.* A**) Diagram showing intranasal vaccination regiment of BALB/c mice with 20 µL volume containing 15 µg *Nm*-nOMV and 15 µg recombinant protein in 10 mM Tris/3% sucrose, pH 7.4 on day 0 and day 21 followed by sacrifice on day 35. **B**) Figure of full spike vaccine candidates with the left column representing SARS-CoV-2 omicron full spike, the middle SARSr-CoV-SHC014 full spike, and the right MERS-CoV full spike. Radar plot showing neutralization profile of pooled sera from mice vaccinated with full spike proteins against S-CoV-2, S-CoV, M-CoV and Sr-CoV VSVpp. Graph keys above the radar plot show the colors corresponding to the dilutions of the sera. **C**) Graphs showing the calculated NT50 values from neutralization assay for full spike candidates. The y-axis represents the sera dilution with the upper limit of deduction (ULOD) at dilution 1:1280 to the corresponding VSVpp shown on the x-axis.

**Table 1:** Vaccine candidates’ overview

| **S. No** | **Candidates** | **Spike protein** | **Species** | **Illustration** | **Dose (µg)** |
|---|---|---|---|---|---|
| 1. | AV-034 | Full | SARS-CoV2 BA.5 |  | 15 |
| 2. | AV-037 | Full | SARSr-CoV- SHC014 |  | 15 |
| 3. | AV-039 | Full | MERS-CoV |  | 15 |
| **Mice immunogenicity study 1** |  |  |  |  |  |
| Combination 1 | AV-002 | Chimeric | SARS-CoV-2 BA.5<br>+<br>SARSr-CoV-SHC014 RBD |  | 7.5 |
|  | AV-014 | Chimeric | MERS-CoV<br>+<br>HKU4 RBD |  | 7.5 |
| Combination 2 | AV-008 | Chimeric | SARSr-CoV-SHC014<br>+<br>Bat-CoV RaTG13 RBD |  | 7.5 |
|  | AV-014 | Chimeric | MERS-CoV<br>+<br>HKU4 RBD |  | 7.5 |
| Combination 3 | AV-014 | Chimeric | MERS-CoV<br>+<br>HKU4 RBD |  | 7.5 |
|  | AV-065 | RBD concatemer (RBD 4) | SARS-CoV 2 BA.5__Bat-CoV RaTG13__SARSr-CoV-SHC014__Bat SARS-CoV Bat WIV |  | 7.5 |
| **Mice immunogenicity study 2** |  |  |  |  |  |
| RBD 1 | AV-085 | RBD concatemer | MERS-CoV__SARS-CoV-2 BA.5__SARS-CoV/2003 |  | 15 |
| RBD 2 | AV-087 | RBD concatemer | SARS-CoV 2 BA.5__Bat-CoV RaTG13__SARS-CoV/2003__ SARSr-CoV-SHC014 __MERS-CoV |  | 15 |
| RBD 3 | AV-088 | RBD concatemer | SARS-CoV 2 BA.5__Bat-CoV RaTG13__SARS-CoV/2003__ SARSr-CoV-SHC014__MERS-CoV__PDF2180 |  | 11.5 |

To attain a broader neutralizing activity against both *Sarbeco* and *Merbeco* S proteins, three *Nm*-nOMV-based combination vaccines were tested intranasally in mice. For comprehensibility AV-002+AV-014 will be addressed as combination 1, AV-008+AV-014 as combination 2 and AV-065+AV-014 as combination 3. Combinations 1 and 2 contained a *sarbeco* and *merbeco* chimeric S protein combinations, and combination 3 included a *Sarbeco* RBD concatemer (AV-065) and a *Merbeco* chimeric S protein. Among the three combinations, combination 3 consisting of a chimeric protein and *sarbeco* RBD concatemer showed the highest breadth in cross neutralization (**Figure 4A**). Combinations 1 and 2 induced neutralizing antibodies with NT50 values ranging from 1:320–1280 for S-CoV-2, Sr-CoV, and M-CoV but failed to induce antibodies against S-CoV (**Figure 4B**). Only combination 3 induced high neutralizing antibody titers (all NT50=1280) (**Figure 4B**) against all four tested VSVpp (**Figure 4A**) and reaching the upper limit of detection (ULOD) with 1:1280. S-CoV-2 and Sr-CoV VSVpp were neutralized by all three combinations despite partial absence of homologous antigens in the vaccines (**Figure 4A**). Previous studies have shown that using RBDs alone as vaccine candidates can lead to increased levels of cross-neutralizing antibodies^31,32^. Hence, we hypothesize that Sr-CoV neutralization in combination 1 and S-CoV-2 neutralization in combination 2 might be attributed to cross-neutralization of antibodies elicited against the SARSr-CoV SHC014 and SARSr-CoV-2 bat-CoV RaTG13 antigens respectively, present in the chimeric S proteins in the combinations (**Figure 4A**). In addition, the presence of Sr-CoV and SARSr-CoV SHC014 as RBDs in combination 3 with RBD concatemer (see **Figure 4A**) might have led to the induction of cross-neutralizing antibodies against S-CoV. The *Sarbeco* RBD concatemer AV-065 (RBD 4) (part of combination 3) was also tested alone and resulted in a comparable antibody neutralization pattern suggesting that the inclusion of RBD-AV-065 along with merbeco chimeric protein in combination 3 increased the breadth of the neutralization antibodies (see **Supplement figure 4A, RBD 4**). However, further studies are required to confirm the observed results.

**Figure 4:**
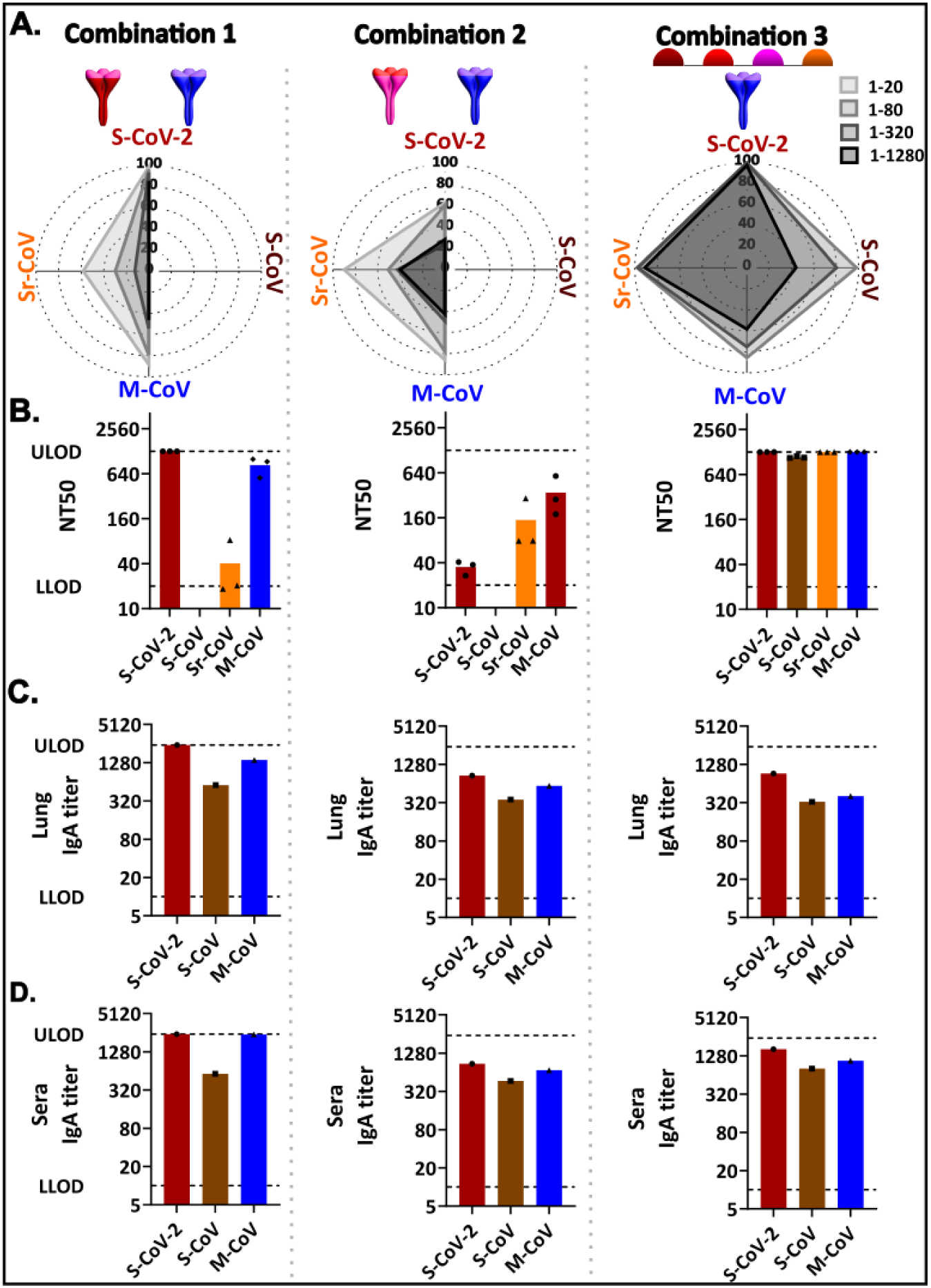
***First mice immunogenicity study – combination groups***. **A**) Figures representing the combination vaccine candidates used in the immunization and the corresponding radar plots showing cross-neutralization profiles of pooled sera from combination vaccine candidates 1,2 and 3 (left to right) against S-CoV-2, S-CoV, M-CoV and Sr-CoV VSVpp with an initial dilution of 1:20 and final dilution of 1:1280 (gray scale light to dark). **B**) Graphs showing the calculated NT50 values from neutralization assay for combination vaccine candidates. The y-axis represents the sera dilution with the upper limit of deduction (ULOD) at dilution 1:1280 to the corresponding VSVpp shown on the x-axis. **C**) IgA ELISA graph with the dilution of pooled lung homogenate on the y-axis with the corresponding spikes used in the ELISA on the x-axis. The bars represent the spike-specific IgA titer in pooled lung homogenates at 0.3 absorbance measured at OD 450nm. **D**) IgA ELISA graph with the dilution of pooled sera on the y-axis with the corresponding spikes used in the ELISA on the x-axis. The bars represent the spike-specific IgA titer in pooled sera at 0.3 absorbance measured at OD 450nm.

Of note, serum samples from the first-mice immunogenicity were also analyzed using micro-NTs (MNT) (**Supplement Figure 2**) using S-CoV-2, S-CoV, and M-CoV viruses. In the full spike candidates, the results from MNTs were analogous to VSVpp-NT assay for both S-CoV-2 and M-CoV viruses. Both the viruses were neutralized by sera from their corresponding full spike vaccinated mice. However, for SARSr-CoV SHC014 full spike, unlike VSVpp-NT assay, there was no cross neutralization observed against S-CoV-2 or S-CoV virus. In the combination of candidates, the neutralization of the viruses was less sensitive compared to the corresponding VSVpp assay. Like VSpp-NT assay, neutralization for S-CoV virus was only observed in vaccine candidate groups that contained AV-065 RBD concatemer. This phenomenon, as previously explained, can be attributed to the presence of Sr-CoV Bat WIV RBD. In conclusion, the obtained results (**Supplement Figure 2**) from the virus-based NTs were overall comparable but less sensitive to the VSVpp assays. Additionally, VSVpp-NT assay could also be performed for bat-borne CoV, for which no full virus isolates were available. Therefore, all subsequent immunogenicity experiments were only tested with VSVpp-NT assays.

Since we vaccinated intranasally, we were next interested in studying markers of the mucosal immunity. For this, both sera and lung homogenates were tested using an IgA and, for comparison, an IgG ELISA. To determine the contributions of the different components of the applied chimeric antigens, we measured immunoglobulins against individual S protein ectodomains comprising S-CoV-2, S-CoV, and M-CoV. All candidates elicited detectable IgA levels in both lung homogenates (**Figure 4C**, range 1:320–2,560) and sera (**Figure 4D**, range 1:640–2,560). The IgG antibody levels were higher, ranging between 1:640–2,560 in lung homogenates (S**upplement Figure 5A**) and all at 1:2,560 for sera (**Supplement Figure 5B**).

Together, we have demonstrated that *Nm*-nOMV-based betacoronavirus chimeric spike and RBD concatemer intranasal vaccinations in mice can induce high IgA levels in lungs and sera as markers for mucosal immunity, high IgG titers, and, most importantly, high levels of broadly neutralizing antibodies. From the NT result data from the first mice immunogenicity study, we identified that the RBD concatemer containing formulation is the most broadly neutralizing design among the candidates tested.

### Second mice immunogenicity study

Results from first mice immunogenicity showed that the concatenated RBD-containing vaccine (combination 3) in combination with the *merbeco* chimeric protein was the most potent and elicited the highest cross-neutralizing antibody titers compared to combination 1 and 2 that contained the same *merbeco* chimeric protein but with 2 different *sarbeco* chimeric proteins (**Figure 4A**). As previously stated, literature has shown that cross-protective antibodies targeting conserved RBD epitopes can neutralize multiple SARS-CoV-2 variants and related betacoronaviruses, including SARS-CoV and WIV-1^12,13^. Hence, to broaden the neutralizing antibody response while simplifying vaccine formulation, we sought to incorporate RBDs from both sarbecoviruses and merbecoviruses into a single multivalent concatemer. Accordingly, recombinant RBD concatemers containing RBD domains from both coronavirus subgenera were designed, expressed, and evaluated for their immunogenicity. These design efforts also focused on optimizing domain borders for protein stability and expression. The tested candidates included AV-085 trimeric (RBD 1), AV-087 pentameric (RBD 2), and AV-088 hexameric (RBD 3). All tested RBD concatemers are illustrated in **Table 1**. The three candidates were tested with the same vaccination protocol as the first mice immunogenicity study. All three newly designed RBD concatemers showed high neutralizing serum titers for all tested VSVpp S protein variants (**Figure 5A-B**) reaching the ULOD of 1:1280 except for M-CoV (**Figure 5B**). Here, only the RBD 3 reached the ULOD for all four tested VSVpp variants. Notably, robust neutralization was achieved for Sr-CoV despite the absence of its RBD in the vaccine candidates, again showing induction of broad-neutralizing antibodies against related *Sarbecoviruses*. Lung homogenates (**Figures 5C**) and sera (**Figure 5D**) showed elevated IgA levels with titers between 1:20–1:320 and 1:160–1:1280, respectively, indicating a detectable lung IgA response. IgG levels ranged between 1:160–1:2560 in lung (**Supplement Figure 6A**) and reached the ULOD of 1:2560 for all VSVpp variants (**Supplement Figure 6B**).

**Figure 5:**
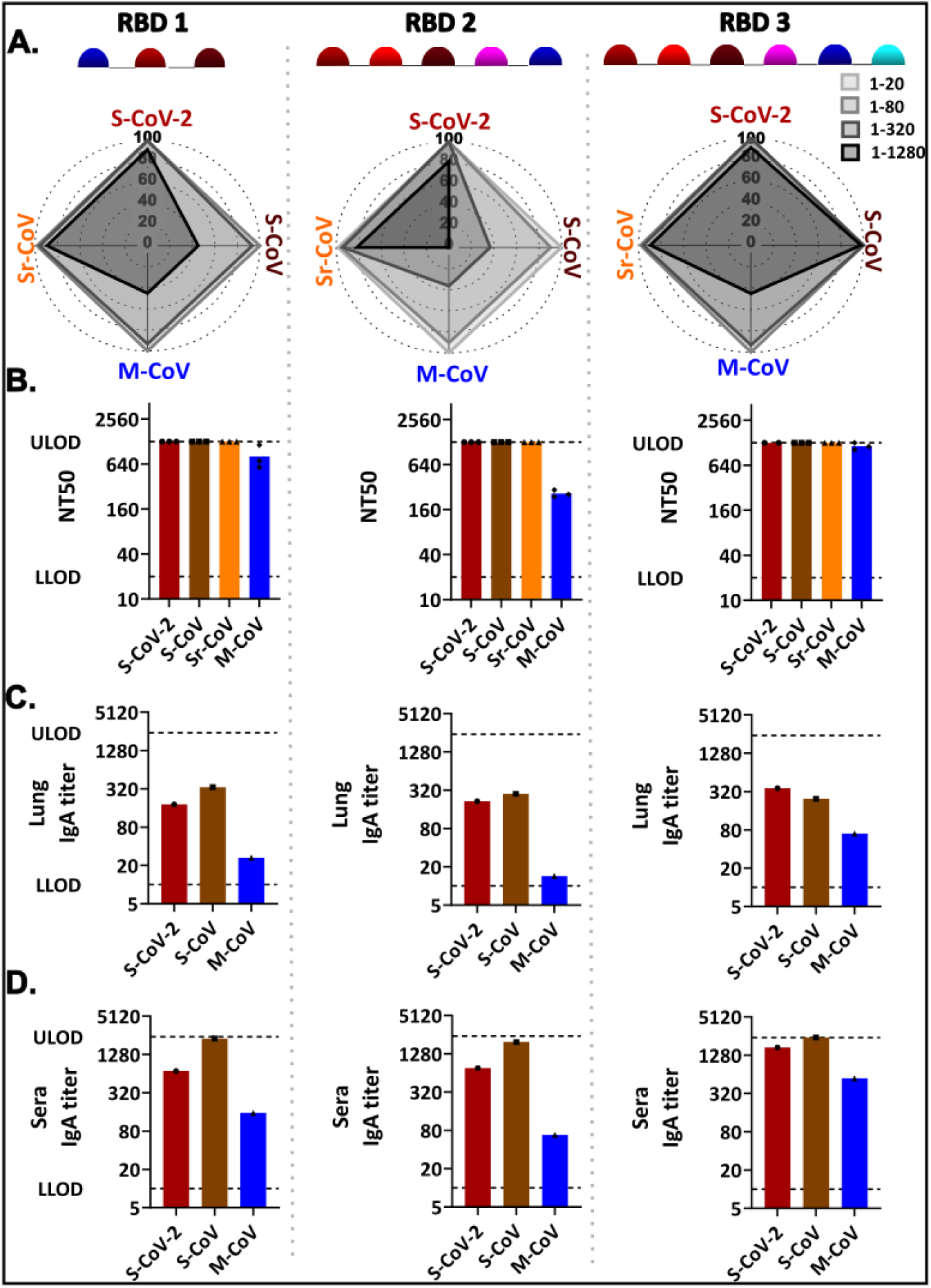
***Second mice immunogenicity study – RBD concatemers.* A**) Radar plots showing cross-neutralization profiles of pooled sera from RBD vaccine candidates 1,2 and 3 (left to right) against S-CoV-2, S-CoV, M-CoV and Sr-CoV VSVpp with an initial dilution of 1:20 and final dilution of 1:1280 (gray scale light to dark). **B**) Graphs showing the calculated NT50 values from neutralization assay for multimeric RBD candidates. The y-axis represents the sera dilution with the upper limit of deduction (ULOD) at dilution 1:1280 to the corresponding VSVpp shown on the x-axis. **C**) IgA ELISA graph with the dilution of pooled lung homogenate on the y-axis with the corresponding spikes used in the ELISA on the x-axis. The bars represent the spike-specific IgA titer in pooled lung homogenates at 0.3 absorbance measured at OD 450nm. D) IgA ELISA graph with the dilution of pooled sera on the y-axis with the corresponding spikes used in the ELISA on the x-axis. The bars represent the spike-specific IgA titer in pooled sera at 0.3 absorbance measured at OD 450nm.

## Discussion

In this manuscript we evaluated an intranasal vaccine strategy that combines a clinically explored *Nm*-nOMV platform with multivalent coronavirus RBD immunogens. Across the antigen formats tested, RBD concatemers provided the greatest breadth of neutralizing antibody responses against both sarbecoviruses and merbecoviruses. In particular, the optimized hexameric RBD concatemer (AV-088) induced neutralizing antibody responses against sarbeco-and merbeco-pseudoviruses, including all highly pathogenic CoVs (S-CoV, S-CoV-2, MERS-CoV), as well as WIV1 whose RBD was absent from the vaccine construct. For the development of a broadly acting betacoronavirus vaccine, literature from SARS-CoV-2 vaccine studies provide no conclusive evidence for the use of full S protein or monomeric RBDs regarding cross-neutralization and protection^33–35^. For a multimeric CoV vaccine, it seems reasonable and beneficial to develop a versatile platform design that uses multimeric proteins for the induction of broadly neutralizing antibodies. Vaccine candidates containing RBD concatemers induced higher cross-neutralizing titers compared to chimeric full S proteins (**Figures 4A and 5A**). We confirmed that the concatenated intranasal RBD-*Nm*-nOMV design is expandable and easily modified for adaptation to newly emerging CoVs (**Table 1**). Our findings support multivalent RBD concatemerization as a modular strategy for broadening antibody responses against genetically diverse CoVs.

CoVs in humans are primarily respiratory pathogens that initiate infection in the mucosal layer of the upper respiratory tract ^36,37^. Consequently, immune responses at the mucosal layer are the first line of defense against viral entry and infection^18^. Secretory IgA in the mucosal layer acts locally and interferes with viral attachment and infection, potentially limiting infection before the virus can disseminate further into the respiratory tract^20,38^. Therefore, compared to systemic vaccination that only induces IgG, intranasal vaccination that induces a combined IgA and IgG response is better suited to protect against CoV infection^17,39,40^. Mucosal immunity is also linked with reduced viral burden and onward transmission in S-CoV-2 animal models^41–43^. Moreover, a study by Dowell and colleagues^44^ reported that mucosal IgA responses against S-CoV-2 and seasonal Betacoronaviruses retained neutralizing activity against emerging Omicron variants, including XBB.1, despite the immune evasion properties of these variants. This suggests that induction of IgA can also contribute to broad protection from other CoV infections. Another recent publication^22^ in which the authors specifically investigated broad-spectrum mucosal CoV vaccine designs demonstrated IgA responses in serum, nasal cavity, and alveolar samples to M-CoV, S-CoV-2, and S-CoV CoV antigens, supporting the feasibility of targeting multiple Betacoronavirus through mucosal vaccination. In our study, multimeric RBD concatemers were combined with *Nm*-nOMVs and delivered by the intranasal route to mice. After immunization, ELISA results of lung homogenates and pooled sera showed production of IgA antibodies against RBDs present in the vaccine candidate (**Figure** 5C). These observations are particularly relevant, as a key objective of a multimeric CoV vaccine is to generate immunity that is not restricted to a single viral strain or variant. Mucosal IgA induced by recognition of RBDs may contribute to this objective because antibodies present at the site of viral entry can potentially recognize conserved epitopes shared among genetically diverse Betacoronaviruses. Thus, the IgA response observed with our vaccine may be considered an important immunological feature supporting its potential as a multi-CoV vaccine.

In the current study, we designed concatenated RBDs -interspaced by short protein linkers - as they provide a distinct, defined antigenic composition and ensure that the selected RBDs are co-delivered to the immune system. This differs from a formulation containing individually expressed RBDs, where differences in protein stability, formulation behavior, biodistribution, or antigen availability could potentially result in unequal exposure to the individual components. One study^31^ demonstrated that nanoparticles displaying RBDs from multiple *Sarbecoviruses* induced better cross-reactive antibody responses compared with nanoparticles displaying only a single S-CoV-2 RBD. Importantly, the mosaic immunogens generated antibodies recognizing heterologous RBDs from *Sarbecoviruses* not represented in the vaccine. This finding was subsequently extended in animal challenge studies, where mosaic-8 RBD nanoparticles induced stronger responses against mismatched *Sarbecoviruses,* increased targeting of conserved RBD epitopes, and provided protection against both S-CoV-2 and S-CoV challenges^31^. Recent studies further support multivalent RBD-based mucosal vaccination as a strategy for broad CoV protection. Intranasally delivered multivalent RBD immunogens combined with STING agonist-based adjuvants induced robust mucosal and systemic immunity and broad protection against diverse Betacoronaviruses in preclinical models^22,45^. A key distinction of the *Nm*-nOMV-based approach is the availability of human clinical safety data. In a recent Phase I trial, intranasal administration of an *Nm*-nOMV-adjuvanted recombinant SARS-CoV-2 spike vaccine was well tolerated and elicited nasal IgA responses^26^. The meningococcal nOMV platform has been previously optimized for human use through deletion of lpxL1, reducing LPS-mediated reactogenicity, and rmpM, enhancing membrane blebbing and OMV production^26,46,47^. Together, this rational attenuation and demonstrated clinical tolerability provide a strong translational basis for *Nm*-nOMV-based mucosal vaccination compared to emerging STING agonist-based platforms, including NanoCF501, for which safety and immunogenicity have thus far been demonstrated only in pre-clinical models.

Furthermore, another study by Walls and colleagues^7^, reported that both mosaic and cocktail RBD Nanoparticle immunogens induced broad neutralizing activity and protected mice against a heterotypic S-CoV challenge, even though the S-CoV RBD was not included in the vaccine. These findings support our strategy of incorporating antigenic diversity into the immunogen to attain a broadly protective multi-CoV vaccine. In addition, this concept has been demonstrated in our study, where we observed antibody induction against S-CoV (**Figure 4A, combination 3**) despite its absence in combination 3 vaccine candidates. Similarly, we have shown neutralizing antibody production against S-CoV related Bat WIV 1 when vaccinated with RBD concatemers containing closely related RBD sequences from other Sr-CoV, namely SARSr-CoV SHC014, in the second immunogenicity study **(Figure 5A**). These observations can only be explained by the production of broad neutralizing antibodies induced by S-CoV and Sr-CoV RBDs present in the vaccine candidates. Taken together, the observed immune responses support that RBD concatemers mixed with *Nm*-nOMVs and delivered via the intranasal route is a promising antigen design strategy for a broad acting intranasal betacoronavirus vaccine.

When designing the RBD concatemers, one of the variables that we considered was the order in which the monomeric RBDs were arranged. As an example, the trimer RBD 5 (**Supplement Figure 4A**) has the same RBD monomers as RBD 1 but with M-CoV monomeric RBD as the last protein in the concatemer. Noticeably, the former elicited lower antibody titers against M-CoV compared to the latter. A comparable trend is observed with the reduced M-CoV neutralizing titers of RBD 2 (**Figure 5A** **RBD 2**), where M-CoV is the terminal RBD in the sequence, in contrast to RBD 1 and RBD 3 (**Figure 5A**). This pattern was not isolated to M-CoV RBD, as a similar effect of reduction in neutralizing titers was observed for S-CoV VSVpp in RBD 1. Here the terminal protein is S-CoV monomeric RBD and shows lower neutralization compared to RBD 5. With two analogous observations, it could be postulated that the position of an RBD within the concatemer could possibly influence its immunogenicity in mice. However, further experiments and designs must be performed to understand the effect of RBD positioning in a vaccine candidate.

To conclude, the S-CoV-2 pandemic brought an increased awareness of Betacoronaviruses that can infect different hosts, leading to a few publications that have described new *Betacoronaviruses* with non-human hosts that can bind human ACE2^48–50^. With these viruses in circulation, it is important to develop an easily administrable and adaptable vaccine with a broad neutralizing potential based on a versatile platform for outbreak preparedness. Moreover, to prevent the spread of respiratory viruses, a vaccine that can elicit mucosal immunity may be particularly advantageous. In the current manuscript, we have described an intranasal vaccine consisting of RBD concatemers combined with *Nm*-nOMVs that can efficiently induce broad neutralizing antibodies, both mucosal and systemic, against multiple *Betacoronavirus* species with a design that has the potential for substitution or addition of RBD monomers for emerging *Betacoronavirus* infections.

The work described is a proof-of-concept study that used a mouse model to study the immune responses generated by B-cells through intranasal immunization. To further prove the feasibility of this vaccine concept, cellular immunity and challenge studies in animal models are required. Furthermore, the current study is focused only on antibody-mediated B-cell response, but we have simultaneously investigated the T-cell immunity where *Nm*-nOMVs incorporate T-cell epitopes separately (manuscript in preparation). The final concept would be a combination of both T-cell components incorporated into *Nm*-nOMVs combined with multimeric RBD concatemers that induce a B-cell response for a broadly protective pan-CoV vaccine.

## Materials and Methods

### Antigen design

Vaccine concepts were designed by combining the S protein receptor-binding domains (RBD) and non-RBD parts of the following viruses belonging to the *Sarbeco*, Merbeco, and *Embeco* subgenera: SARS-CoV-2 Omicron BA.5 (USQ23808), RaTG13 (QHR63300), SHC014 (QJE50589), SARS-like CoV WIV1 (AGZ48828), HKU-39849 (AY278491), MERS-CoV (YP_007188579), PDF-2180 (YP_009361857), Bat-CoV HKU4 (YP_001039953), HCoV-OC43 (UPG19289/UTM04202.1) and HCoV-HKU1 (ADN03339/Q0ZME7) (**Figures** 1A and 1B).

Recombinant proteins were produced by ExcellGene (Monthey, Switzerland) on CHOExpress cells. Ni Sepharose excel resin (Cytiva) was used to purify the His-tagged proteins, which was followed by size exclusion purification using Superdex 200 pg (Cytiva). Proteins were eluted in 10 mM Tris, 3% sucrose, pH 7.4. Protein concentrations were determined using UV absorbance at 280 nm and purities by HPLC-SEC ^51,52^ (**Supplement Table 2**).

### Nm-nOMV production and characterization

*N. meningitidis* native OMVs (AdJane Biosciences’) were produced as described previously^21^. The sterile filtered *Nm*-nOMVs were stored in 10 mM Tris/3% sucrose, pH 7.4, at 2-8°C. Size distribution of the vesicles was determined using dynamic light scattering (DLS). Mass spectrometry was used to analyze the protein composition, while the total protein concentration was determined by the Peterson assay.

### Immunofluorescence assay (IFA)

Initially, endpoint antibody titers against seven different CoV S proteins were determined using a previously described recombinant S protein-based immunofluorescence assay for selection of human reference sera^27^. Sera from twenty healthcare workers who had previously tested virus RNA-positive for the common cold HCoV-NL63, 229E, OC43, and HKU1 were collected at two time points. The first collection time point was before SARS-CoV-2 vaccination (V1). The second time point was four weeks after a SARS-CoV-2 vaccination (Comirnaty, BionTech/Pfizer) (V3). All V1 and V3 sera were then characterized by IFA that is based on cells expressing recombinant S proteins^27^.

### Enzyme linked immunosorbent assay (ELISA) mAb and IgA/IgG

96-well plates (NUNC^TM^ MaxiSorp) were coated with 100 ng of recombinant purified S protein ectodomain per well and incubated at room temperature for 3 hours. The coated plates were washed three times with PBS + 0.05% Tween-20 (PBS-T) wash buffer and blocked with 5% milk powder in PBS-T (blocking buffer) and stored at -20°C. To test reactivity of animal sera, a 4x serial dilution of the sera (in blocking buffer, starting from 1:100) was prepared. 100 µl of the dilution series were then transferred to the pre-coated plates and incubated at room temperature for one hour. For mAb, 100 µl of the diluted antibodies (0.0001ug to 10ug) were added to the precoated plates and incubated at room temperature for 1 hour. Plates were washed, and 100 µl/well 1-8000 anti-mouse-IgG-HRP or 1-4000 anti-mouse-IgA-HRP (Southern Biotech, Birmingham, USA) for mice sera and 1-5000 dilutions of goat anti-human-HRP (Invitrogen, A18805) for mAbs was incubated at 37°C for 60 minutes. The wash step was repeated, and 100 µl of TMB Super Slow One Component HRP Microwell substrate (BioFX) was added and incubated in the dark for 5 minutes (IgG) or 10 minutes (IgA). Finally, the reaction was stopped by the addition of 100 µl of 12.5% H_2_SO_4_ and the resulting endpoint absorbance was quantified by using a BioTek 800 TS Microplate Reader at a 450 nm wavelength.

### Enzyme linked immunosorbent assay (ELISA) human coronavirus-reactive sera

For each human serum, 96 well polystyrene plates (PierceTM Thermofischer) were coated with 150 ng of recombinant protein per well in triplicates and incubated at 4°C overnight. The coated plates were washed three times with 1XTBS-Tween (TBS-T) wash buffer (PierceTM Thermofisher) the following day and blocked with 2% BSA Pan-Biotech in TBS-T at ambient temperature for one hour. Succeeding the blocking, 100 µl of pre-defined sera diluted with blocking buffer was added and incubated at 37°C for one hour. The dilution series of the sera consisted of 2-fold dilutions starting at 1:200 to 1:6400. Plates were washed and 100 µl/well Anti-human-IgG (EUROIMMUN AG, Lübeck, Germany) secondary antibody enzyme mix was added followed by incubation at 37°C for 30 minutes. The wash step was repeated and 100 µl of One-step TMB substrate (EUROIMMUN AG) was added and incubated in the dark for 30 minutes. Finally, the reaction was stopped by addition of 100 µl of stop solution (EUROIMMUN AG) and the resulting endpoint absorbance was quantified by using BioTek Synergy HTX Multimode Reader at 450 nm wavelength.

### Immunogenicity study

BALB/c mice (OlaHSD; Envigo, female, 8–9 weeks old at day 0), 10 animals per group, were immunized intranasally (i.n.) on days 0 and 21 (**Figure 3A**). Each dose consisted of a 20 µL volume containing 15 µg *Nm*-nOMV and 15 µg recombinant protein in 10 mM Tris/3% sucrose, pH 7.4, mixed just prior to immunization. The inoculum was divided over both nostrils using a pipette. On days 0, 21, and 35, blood samples were collected for determination of a neutralizing antibody response in the sera. On day 35 animals were euthanized under anesthesia (isoflurane/O2) by exsanguination, after which lungs were collected.

### Microneutralization test

Sera from the first mouse immunogenicity study was tested for the presence of neutralizing antibodies using a microneutralization test with MERS-CoV, SARS-CoV-1 or SARS-CoV-2 viruses, at a concentration of 100 spots/well. Viruses were incubated with twofold dilutions of the animal sera for 1 hour at RT. Next, mixtures were added to Vero E6 cells and incubated at 37°C for 1 hour. Inocula were removed from the cells and replaced by an infection medium containing 1.6% CMC after which plates were incubated at 37°C for ∼20 hours. After this incubation, the cells were fixed by formalin and subsequently permeabilized by ethanol. Cells were immuno-stained by using a viral nucleocapsid targeting murine monoclonal antibody, followed by a secondary anti-mouse IgG peroxidase conjugate. Upon addition of TrueBlue substrate, a blue precipitate formed on nucleocapsid-positive cells. Images were acquired of all wells by a CTL ImmunoSpot analyzer, equipped with software to quantitate the nucleocapsid-positive and, thus, infected cells. MN50, MN80, and (**Supplement Figure 1B**) MN90 values were calculated according to the method described by Zielinska and colleagues^53^.

### VSV-based pseudoparticle production

Human codon-optimized *Betacoronavirus* S protein sequences lacking the C-terminal last 19 aa that contain an ER retention signal were cloned into pCG1 vector using In-Fusion® Snap Assembly (TAKARA, Japan). Spike carrying Vesicular Stomatitis Virus pseudoparticles (VSVpp) with a GFP reporter were generated in HEK293T cells, as previously described^54^. Synthesized particles were titrated on Vero E6 TMPRSS2 cells, aspirated and, fixed with ROTI®Histofix (Carl Roth) for 15 minutes, succeeded by 5 minutes of staining with DAPI (1 to 500 dilution of 1 mg/mL, Art.-Nr. 3105.2, D9542-1MG) for cell confluency control. The resulting foci-forming units (FFUs) were imaged using the Autoimmun Diagnostika V Spot Spectrum. The FFUs with intensity ≥30 was counted to determine the titer of the pseudoparticles stocks.

### Neutralization test (NT) with Pseudoparticles

Diluted sera (1:10–1:1280) or the positive control mAb 7.7G6 (1:20,000–1:400,000)^55^ were mixed with 200 foci-forming units (FFUs) of VSVpp and incubated at 37°C for 30 minutes. Following incubation, the mixture was used to infect Vero E6 TMPRSS2 cells in triplicates in 96 wells that were plated 24 hours prior to the experimental setup. The plates were incubated for 18 hours at 37°C. Following the final incubation, cells were fixed and imaged, and the FFUs were counted as described above, and the neutralization value was calculated.

## Supplement Figures and Tables

### Ethics statement

Animal studies were reviewed and approved by Instantie voor Dierenwelzijn Utrecht (IvD Utrecht, Animal Welfare Body Utrecht).

## Acknowledgements

This study was funded by Coalition for Epidemic Preparedness Innovations (CEPI), grant number: PRJ-6794. The funder played no role in study design, data collection, analysis and interpretation of data, or the writing of this manuscript. We want to thank Nicolas Hienemann for his technical support. We would like to acknowledge Dr. Roberto Cattaneo for sharing the spike expression plasmids pCG1. We also want to acknowledge Maria Grazia Pau (CSO, AdJane Biosciences) for her review of this manuscript.

## Author contributions

KH and RH developed the methodology, performed the investigations and formal analysis, and contributed to writing the original draft. KH additionally prepared the figures. WB contributed to writing the original draft, performed formal analysis and was responsible for project administration and provision of resources. JL, NK and FR performed the experimental investigations. DO, CD, MM, BB and CK conceptualised the study. MM, BB and CK were additionally supervised the research activities and reviewed and edited the manuscript. DO and CK acquired funding for the research.

---

**Supplement Figure 1:**
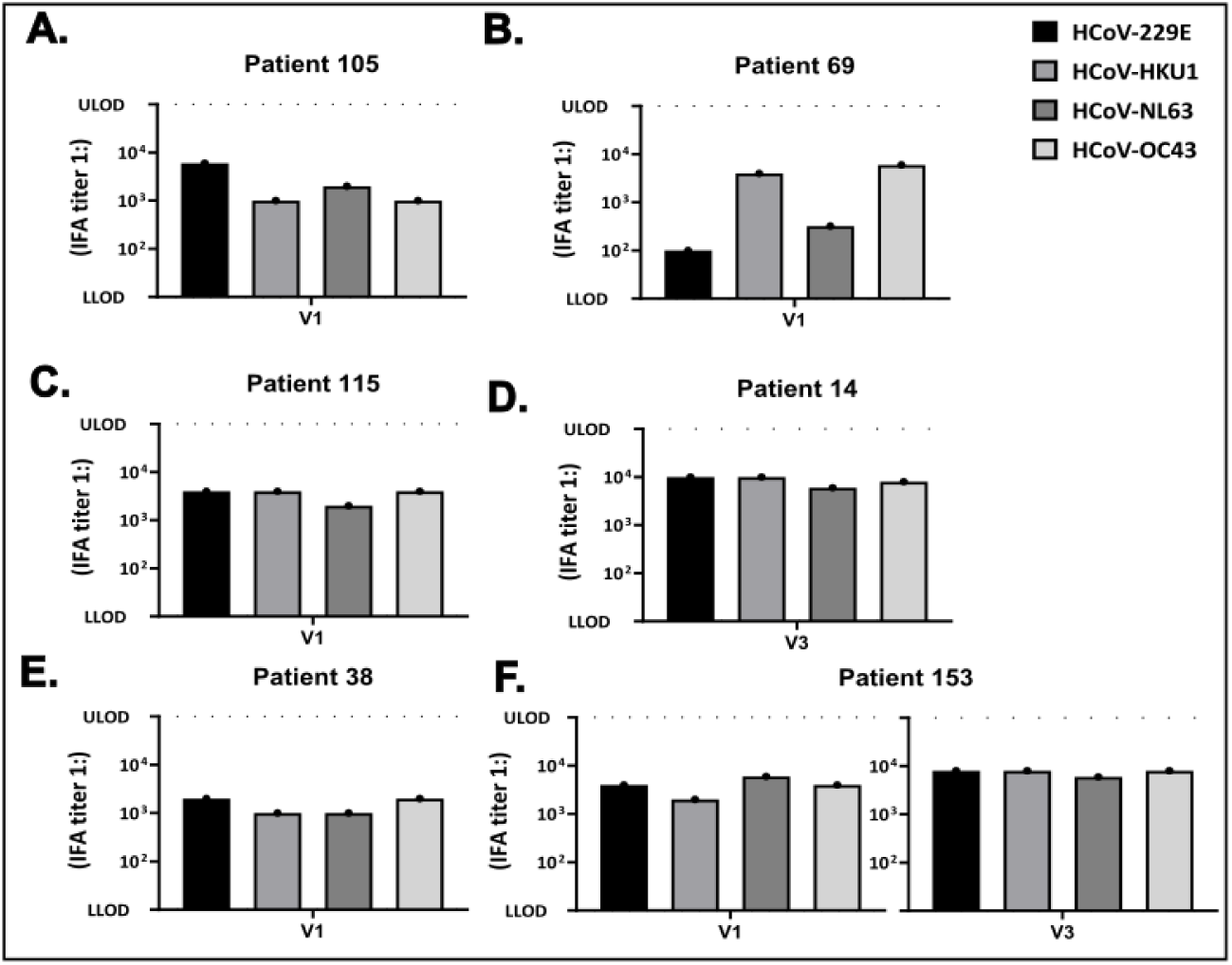
***Serum characterization.*** Graphs A to F showing IFA end point titer of a subset of sera collected from health care workers before (V1) and after (V3) SARS-CoV-2 vaccination. TY-axis showing the endpoint titer against the different HCoV spikes on the x-axis.

**Supplement Figure 2:**
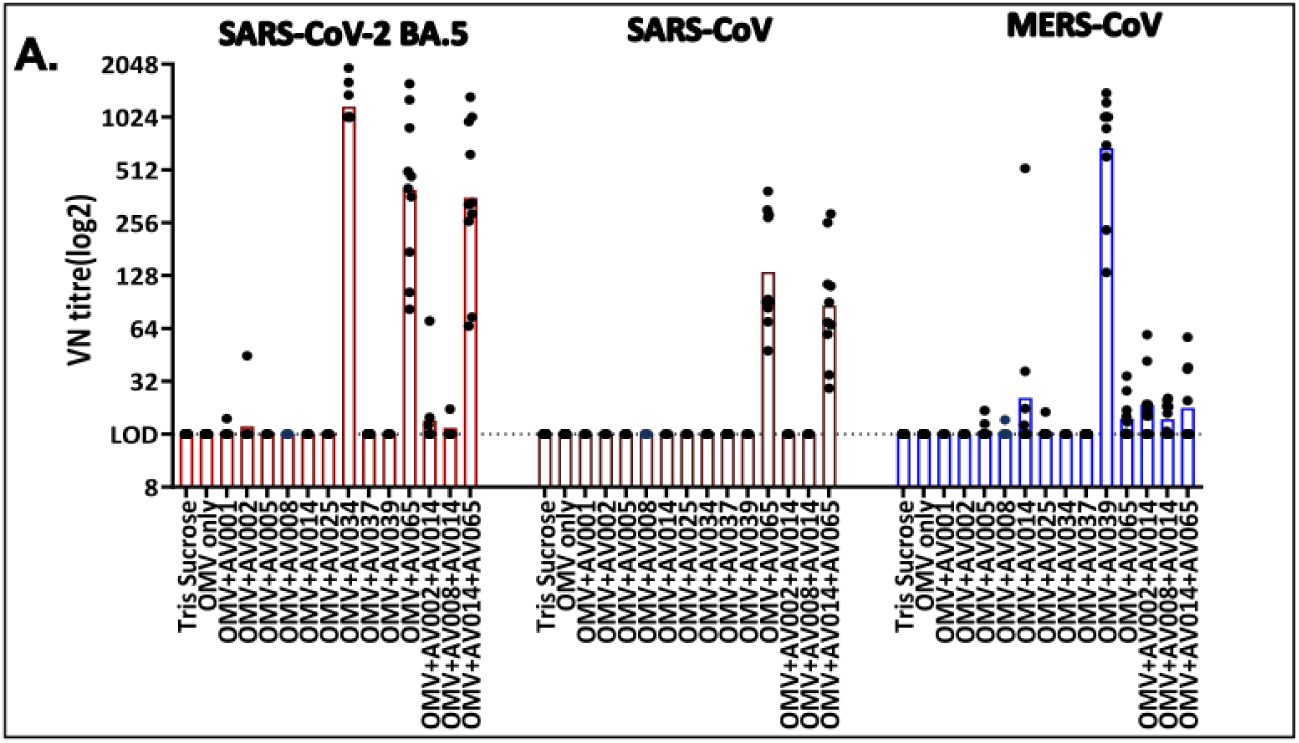
***Microneutralization test*. A)** Graph showing NT90 values of virus microneutralization assay of mice sera from the first immunogenicity study with vaccine candidates on x-axis and serum dilutions on y-axis. The NT90 values are grouped according to the viruses that the sera were tested against.

**Supplement Figure 3:**
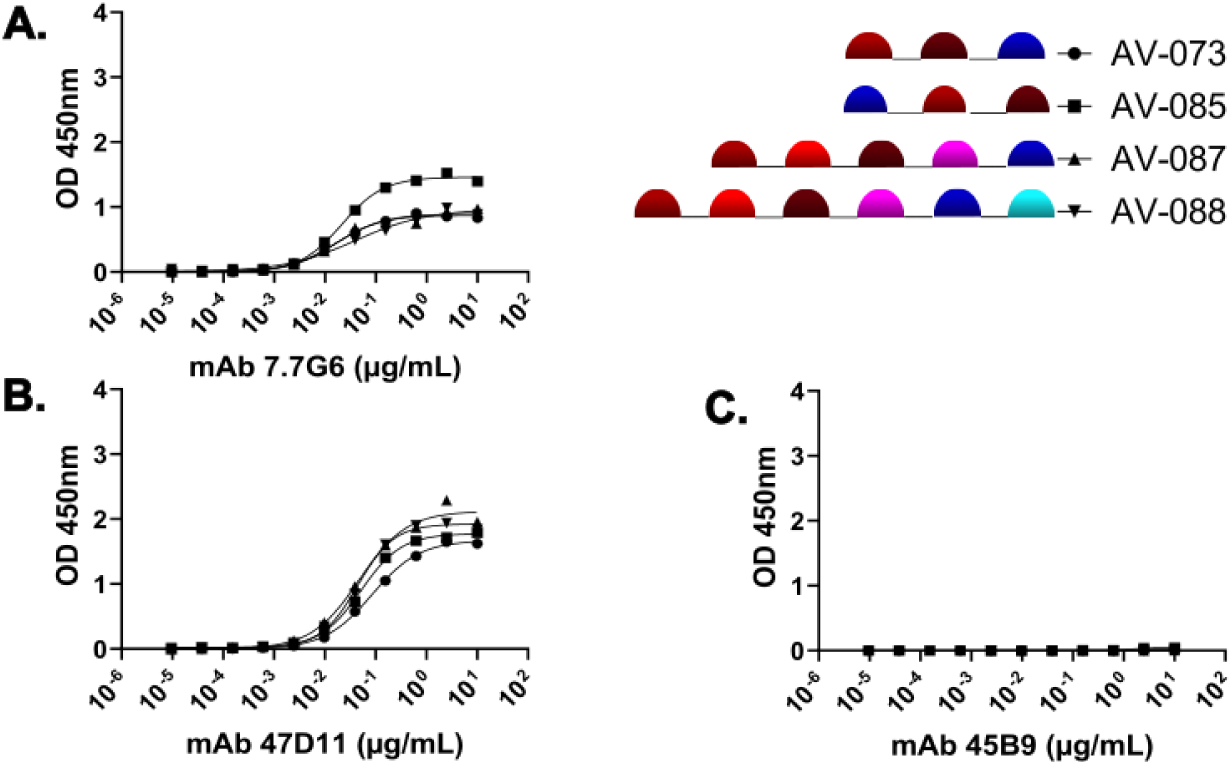
***Antigenic profiling using coronavirus spike-specific monoclonal antibodies***. **A**) Graph showing results of RBD vaccine candidate screen ELISA assay with mAb 47D11 (conformational Ab against SARS-CoV-2 S1B/RBD region). The x-axis represents dilution of antibodies, and the y-axis represents OD at 450 nm. **B**) Graph showing results of RBD vaccine candidate screen ELISA assay with mAb 7.7G3 (conformational Ab against MERS-CoV S1B/RBD region). **C**) Graph showing results of RBD vaccine candidate screen ELISA assay with mAb 45B9 (conformational Ab against HCoV-OC43 strains, BCoV, and ECoV S1B/RBD region).

**Supplement Figure 4:**
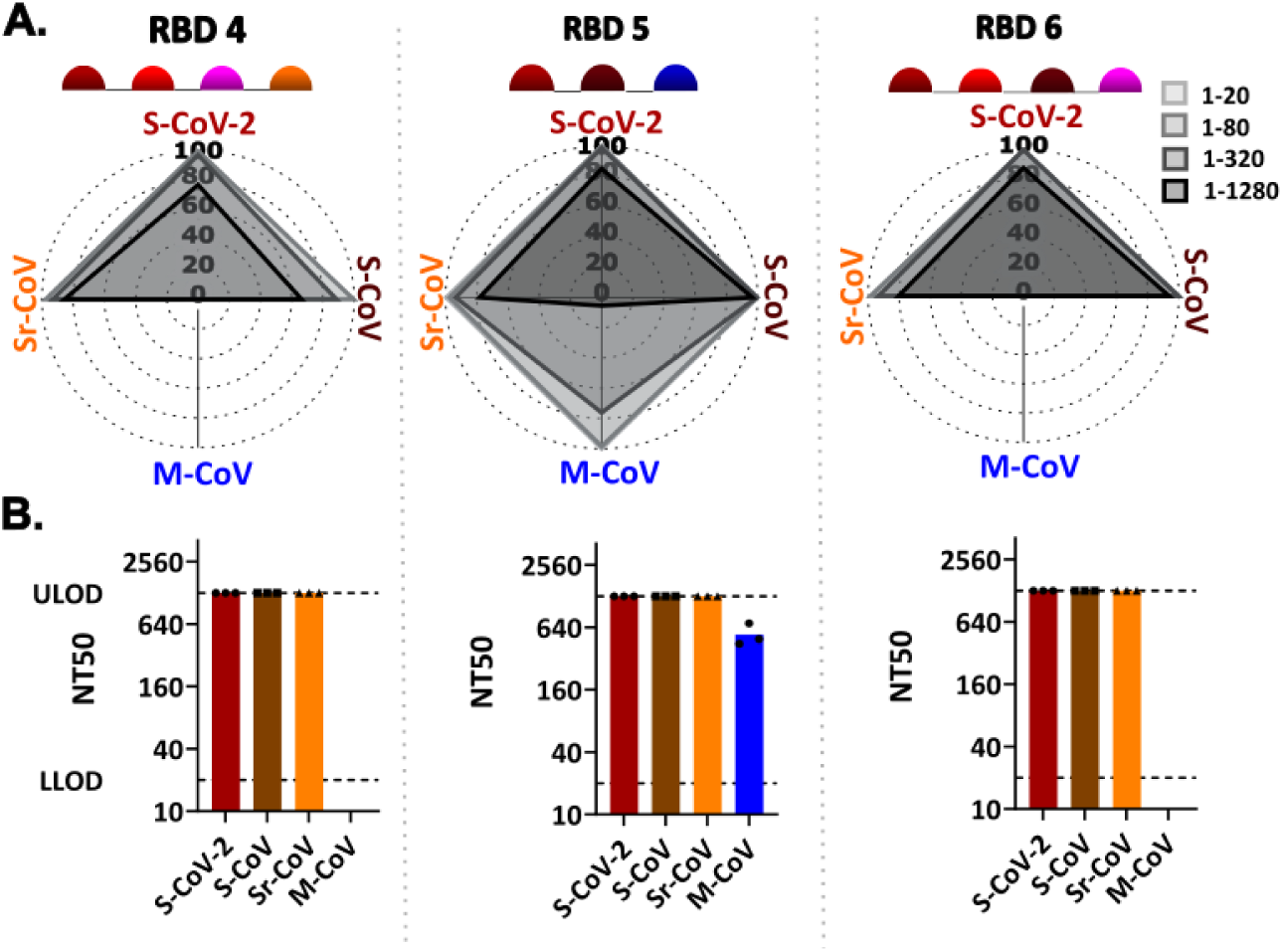
***Second mice immunogenicity study.* A**) Radar plots showing cross-neutralization profiles of pooled sera from RBD vaccine candidates 4, 5 and 6 (left to right) against S-CoV-2, S-CoV, M-CoV and Sr-CoV VSVpp with an initial dilution of 1:20 and final dilution of 1:1280 (gray scale light to dark). **B**) Graphs showing the calculated NT50 values for the first mice immunogenicity candidates on the y-axis (ULOD 1 to 1280) to the corresponding VSVpp shown on the x-axis.

**Supplement Figure 5:**
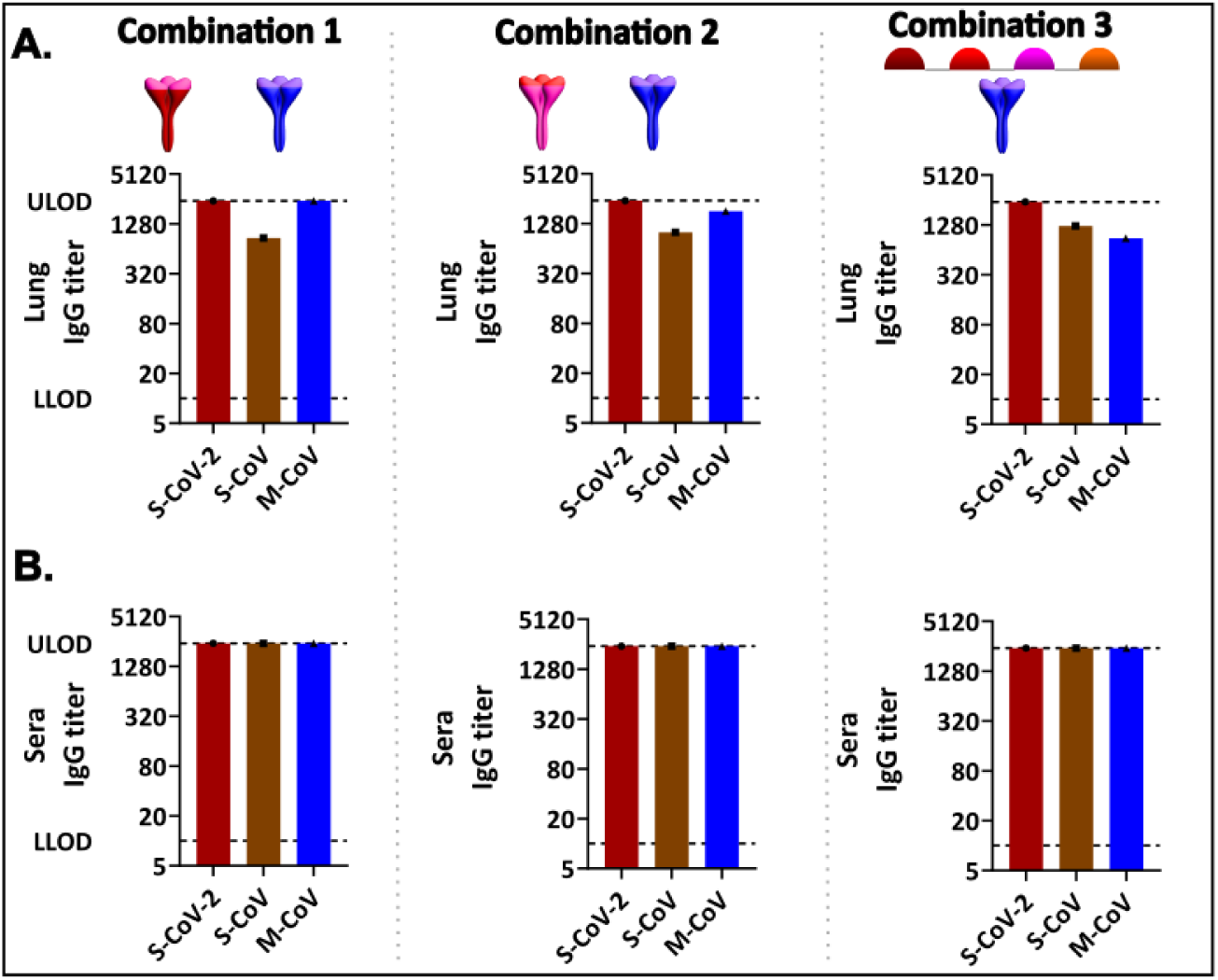
***First mice immunogenicity study with combination candidates.* A**) IgG ELISA graph for combination candidates 1, 2 and 3 (columns left to right), with the dilution of pooled lung homogenates on the y-axis with the corresponding spikes used in the ELISA on the x-axis. The bars represent the spike-specific IgG titer in pooled lung homogenates at 0.3 absorbance measured at OD 450nm. **B**) IgG ELISA graph with the dilution of pooled sera on the y-axis with the corresponding spikes used in the ELISA on the x-axis. The bars represent the spike-specific IgG titer in pooled sera at 0.3 absorbance measured at OD 450nm.

**Supplement Figure 6:**
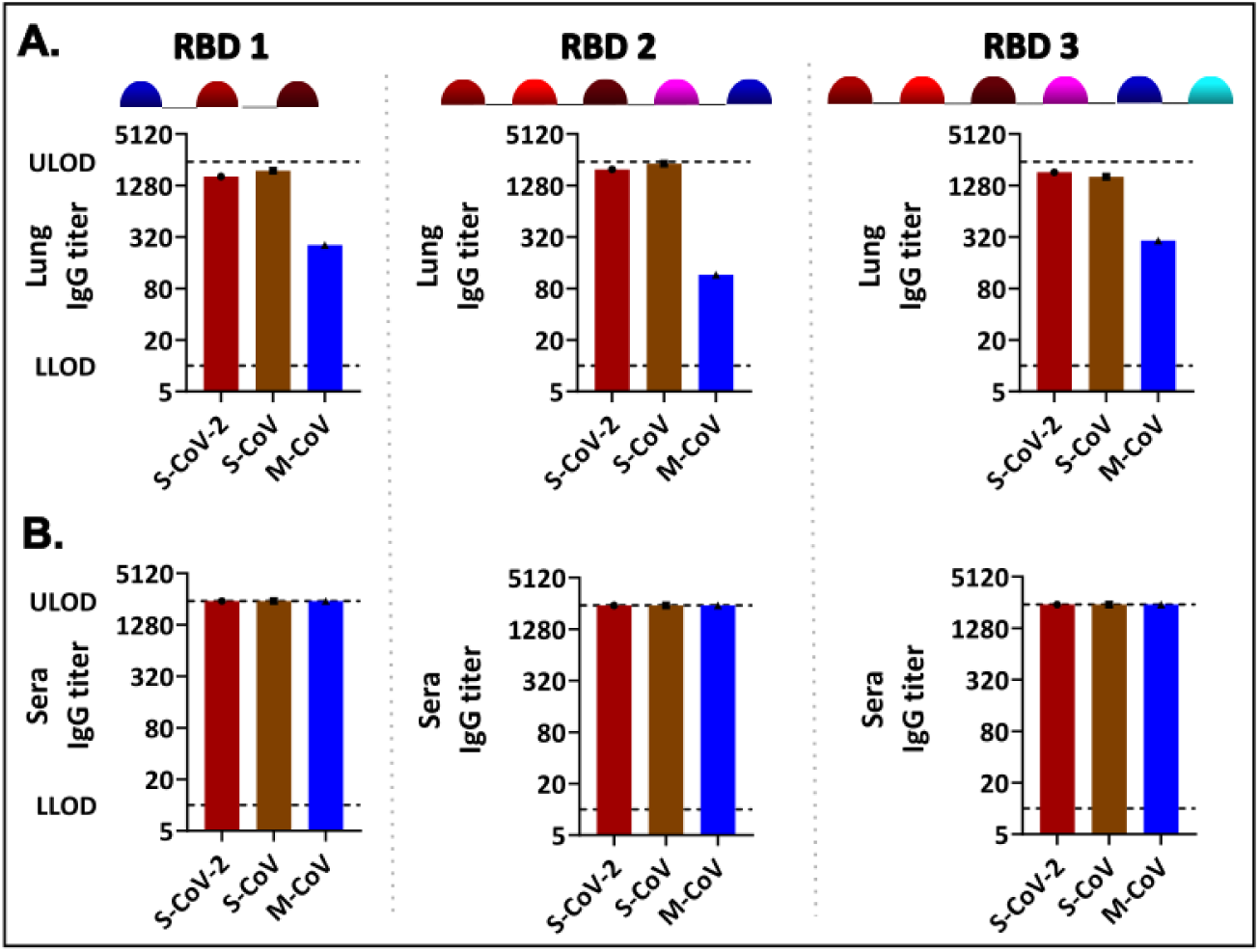
***Second mice immunogenicity study with multimeric RBD concatemers.* A**) IgG ELISA graph for multimeric RBD candidates 1, 2 and 3 (columns left to right), with the dilution of pooled lung homogenate on the y-axis with the corresponding spikes used in the ELISA on the x-axis. The bars represent the spike-specific IgG titer in pooled lung homogenates at 0.3 absorbance measured at OD 450nm. **B**) IgG ELISA graph with the dilution of pooled sera on the y-axis with the corresponding spikes used in the ELISA on the x-axis. The bars represent the spike-specific IgG titer in pooled sera at 0.3 absorbance measured at OD 450nm.

**Supplement Table 1:**
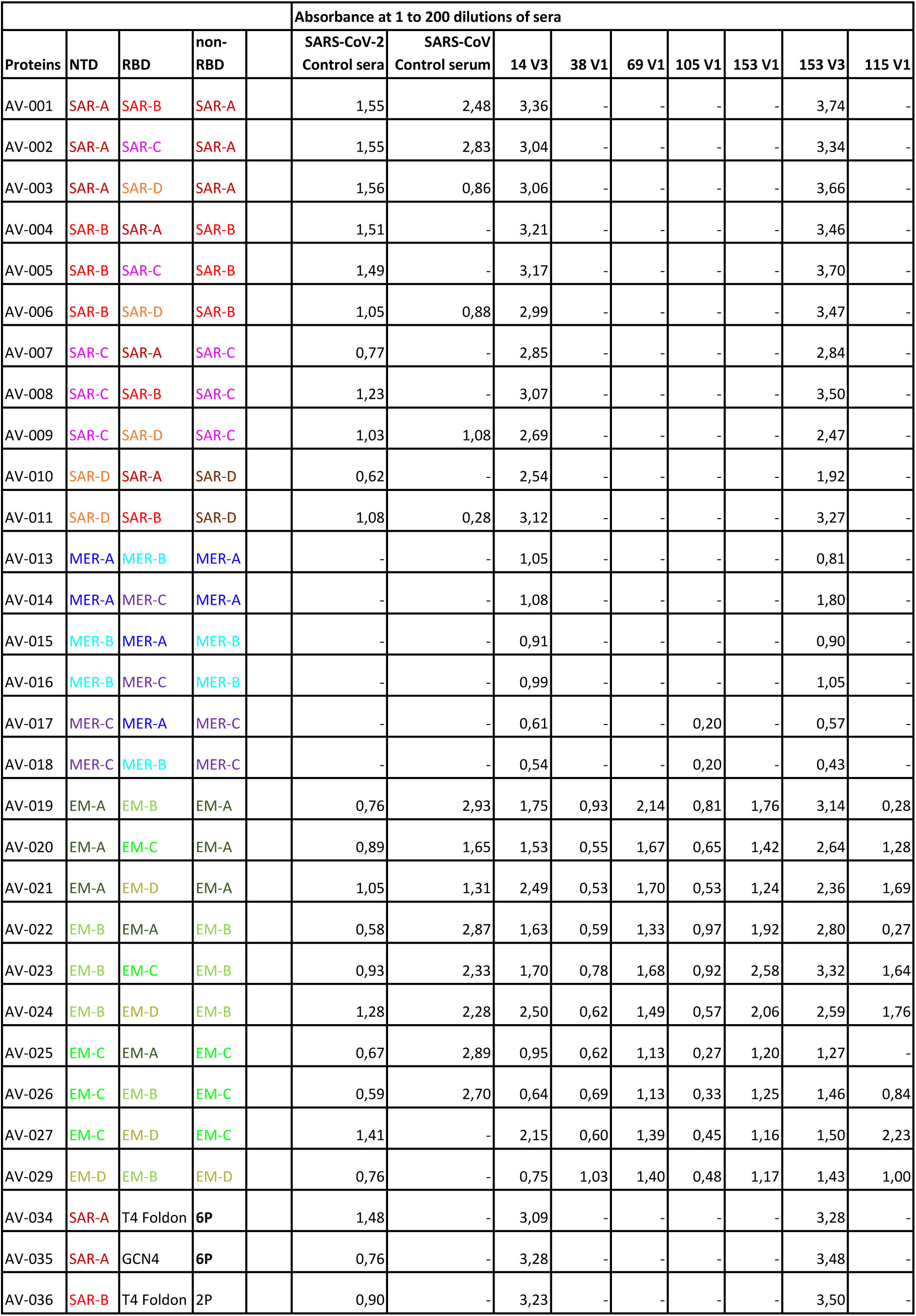
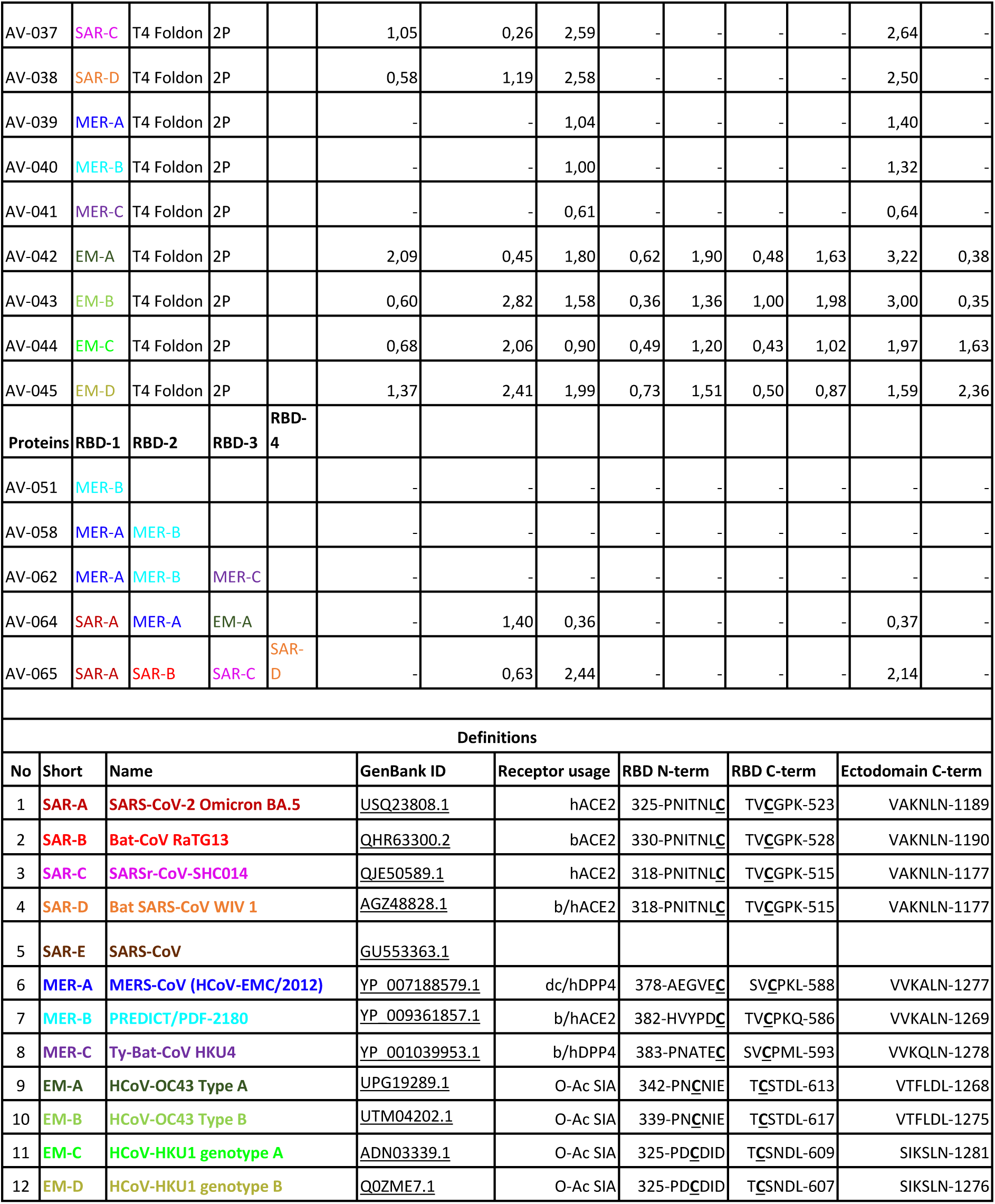
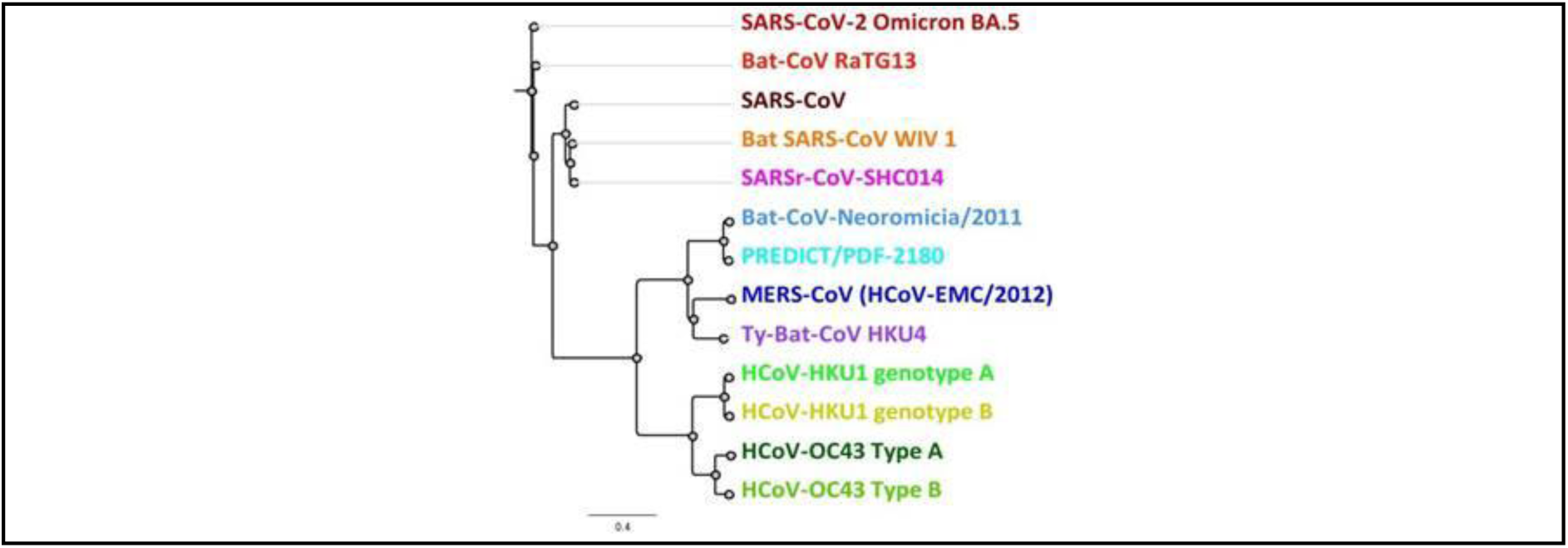
Antigenic profiling of vaccine candidates using human coronavirus-reactive sera.

**Supplement Table 2:** Vaccine candidate quality attributes.

| Proteins | NTD | RBD | non-RBD |  |  |  | Conc. (mg/mL) | Protein Amount (mg) | HMW (%) | Main Purity (%) | LMW (%) | EU/mL | Mouse study |
|---|---|---|---|---|---|---|---|---|---|---|---|---|---|
| AV-001 | SAR-A | SAR-B | SAR-A |  |  |  | 1,05 | 13,7 | 13,12 | 78,96 | 7,93 | < 0.01 | 1 |
| AV-002 | SAR-A | SAR-C | SAR-A |  |  |  | 1,17 | 24,6 | 8,1 | 82,9 | 8,9 | < 0.01 |  |
| AV-005 | SAR-B | SAR-C | SAR-B |  |  |  | 0,78 | 7,1 | 12,99 | 79,8 | 7,21 | < 0.01 |  |
| AV-008 | SAR-C | SAR-B | SAR-C |  |  |  | 0,83 | 22,5 | 10,2 | 80,9 | 8,9 | 0,39 |  |
| AV-014 | MER-A | MER-C | MER-A |  |  |  | 1,05 | 6,3 | 21,7 | 70,5 | 7,8 | < 0.01 |  |
| AV-025 | EM-C | EM-A | EM-C |  |  |  | 1,02 | 18,4 | 13,7 | 78,7 | 7,6 | 1,45 |  |
| AV-034 | SAR-A |  |  |  |  |  | 1,17 | 31,5 | 8,1 | 80,9 | 11,0 | < 0.01 |  |
| AV-037 | SAR-C |  |  |  |  |  | 1,13 | 12,5 | 10,1 | 85,0 | 5,0 | 1,39 |  |
| AV-039 | MER-A |  |  |  |  |  | 0,92 | 15,6 | 15,7 | 76,6 | 7,7 | < 0.01 |  |
| **Proteins** | **RBD-1** | **RBD-2** | **RBD-3** | **RBD-4** | **RBD-5** | **RBD-6** |  |  |  |  |  |  |  |
| AV-065 | SAR-A | SAR-B | SAR-C | SAR-D |  |  | 1,43 | 30,1 | 1,3 | 94,6 | 4,1 | 2,04 | 1 & 2 |
| AV-067 | SAR-A |  |  |  |  |  | 2,01 | 10,3 | 1,0 | 98,1 | 1,0 | <0.10 | 2 |
| AV-068 | SAR-E |  |  |  |  |  | 2,26 | 13,7 | 0,3 | 99,4 | 0,3 | <0.10 |  |
| AV-073 | SAR-A | SAR-E | MER-A |  |  |  | 1,03 | 6,7 | 27,5 | 61,9 | 10,6 | 0,13 |  |
| AV-081 | SAR-A | SAR-B | SAR-E | SAR-C |  |  | 1,07 | 5,7 | 27,4 | 59,9 | 12,7 | 0,25 |  |
| AV-085 | MER-A | SAR-A | SAR-E |  |  |  | 0,87 | 2,1 | 36,3 | 51,7 | 11,9 | <0.10 |  |
| AV-087 | SAR-A | SAR-B | SAR-E | SAR-C | MER-A |  | 1,11 | 2,6 | 23,9 | 59,2 | 17,0 | 0,27 |  |
| AV-088 | SAR-A | SAR-B | SAR-E | SAR-C | MER-A | MER-B | 0,67 | 1,6 | 21,7 | 54,2 | 24,1 | 0,20 |  |

**Supplement Table 3:** Monoclonal antibody specificity

| **Monoclonal antibody** | **Targeted domain** | **Epitope** | **Binding breadth** | **Reference (PMID)** |
|---|---|---|---|---|
| 47D11 | S1B (=RBD) | Conformational | SARS-CoV-2 (but not Omicron), WIV16 | 32366817 |
| 7.7G3 | S1B (=RBD) | Conformational | MERS | 30938227 |
| 45B9 | S1B (=RBD) | Conformational | HCoV-OC43 strains, BCoV, ECoV | 35614127 |

## References

1 Cui, J., Li, F. & Shi, Z. L. Origin and evolution of pathogenic coronaviruses. Nat Rev Microbiol 17, 181–192 (2019). 10.1038/s41579-018-0118-9

2 Graham, R. L., Donaldson, E. F. & Baric, R. S. A decade after SARS: strategies for controlling emerging coronaviruses. Nat Rev Microbiol 11, 836–848 (2013). 10.1038/nrmicro3143

3 Frutos, R., Serra-Cobo, J., Pinault, L., Lopez Roig, M. & Devaux, C. A. Emergence of Bat-Related Betacoronaviruses: Hazard and Risks. Front Microbiol 12, 591535 (2021). 10.3389/fmicb.2021.591535

4 Ruiz-Aravena, M. et al. Ecology, evolution and spillover of coronaviruses from bats. Nat Rev Microbiol 20, 299–314 (2022). 10.1038/s41579-021-00652-2

5 Krammer, F. SARS-CoV-2 vaccines in development. Nature 586, 516–527 (2020). 10.1038/s41586-020-2798-3

6 Bosch, B. J., van der Zee, R., de Haan, C. A. & Rottier, P. J. The coronavirus spike protein is a class I virus fusion protein: structural and functional characterization of the fusion core complex. J Virol 77, 8801–8811 (2003). 10.1128/jvi.77.16.8801-8811.2003

7 Walls, A. C. et al. Elicitation of broadly protective sarbecovirus immunity by receptor-binding domain nanoparticle vaccines. Cell 184, 5432–5447 e5416 (2021). 10.1016/j.cell.2021.09.015

8 Barnes, C. O. et al. SARS-CoV-2 neutralizing antibody structures inform therapeutic strategies. Nature 588, 682–687 (2020). 10.1038/s41586-020-2852-1

9 Wrapp, D. et al. Cryo-EM structure of the 2019-nCoV spike in the prefusion conformation. Science 367, 1260–1263 (2020). 10.1126/science.abb2507

10 Cao, Y. et al. Imprinted SARS-CoV-2 humoral immunity induces convergent Omicron RBD evolution. Nature 614, 521–529 (2023). 10.1038/s41586-022-05644-7

11 Planas, D. et al. Considerable escape of SARS-CoV-2 Omicron to antibody neutralization. Nature 602, 671–675 (2022). 10.1038/s41586-021-04389-z

12 Chen, Y. et al. Broadly neutralizing antibodies to SARS-CoV-2 and other human coronaviruses. Nat Rev Immunol 23, 189–199 (2023). 10.1038/s41577-022-00784-3

13 Tortorici, M. A. et al. Broad sarbecovirus neutralization by a human monoclonal antibody. Nature 597, 103–108 (2021). 10.1038/s41586-021-03817-4

14 Cohen, A. A. et al. Mosaic nanoparticles elicit cross-reactive immune responses to zoonotic coronaviruses in mice. Science 371, 735–741 (2021). 10.1126/science.abf6840

15 Singanayagam, A. et al. Community transmission and viral load kinetics of the SARS-CoV-2 delta (B.1.617.2) variant in vaccinated and unvaccinated individuals in the UK: a prospective, longitudinal, cohort study. Lancet Infect Dis 22, 183–195 (2022). 10.1016/S1473-3099(21)00648-4

16 Eyre, D. W. et al. Effect of Covid-19 Vaccination on Transmission of Alpha and Delta Variants. N Engl J Med 386, 744–756 (2022). 10.1056/NEJMoa2116597

17 Hassan, A. O. et al. A Single-Dose Intranasal ChAd Vaccine Protects Upper and Lower Respiratory Tracts against SARS-CoV-2. Cell 183, 169–184 e113 (2020). 10.1016/j.cell.2020.08.026

18 Russell, M. W., Moldoveanu, Z., Ogra, P. L. & Mestecky, J. Mucosal Immunity in COVID-19: A Neglected but Critical Aspect of SARS-CoV-2 Infection. Front Immunol 11, 611337 (2020). 10.3389/fimmu.2020.611337

19 Brandtzaeg, P. Secretory IgA: Designed for Anti-Microbial Defense. Front Immunol 4, 222 (2013). 10.3389/fimmu.2013.00222

20 Corthesy, B. Multi-faceted functions of secretory IgA at mucosal surfaces. Front Immunol 4, 185 (2013). 10.3389/fimmu.2013.00185

21 van der Ley, P. A., Zariri, A., van Riet, E., Oosterhoff, D. & Kruiswijk, C. P. An Intranasal OMV-Based Vaccine Induces High Mucosal and Systemic Protecting Immunity Against a SARS-CoV-2 Infection. Front Immunol 12, 781280 (2021). 10.3389/fimmu.2021.781280

22 Wu, Y. et al. Universal broad-spectrum mucosal vaccine design for human coronaviruses inspired by artificial antibodies. NPJ Vaccines 11, 55 (2026). 10.1038/s41541-026-01375-2

23 Zhou, J. et al. Monoclonal antibodies target conserved S1(B) epitopes across deltacoronaviruses to inhibit porcine deltacoronavirus infection. mBio 17, e0174226 (2026). 10.1128/mbio.01742-26

24 Etchart, N. et al. Intranasal immunisation with inactivated RSV and bacterial adjuvants induces mucosal protection and abrogates eosinophilia upon challenge. Eur J Immunol 36, 1136–1144 (2006). 10.1002/eji.200535493

25 Sardinas, G., Reddin, K., Pajon, R. & Gorringe, A. Outer membrane vesicles of Neisseria lactamica as a potential mucosal adjuvant. Vaccine 24, 206–214 (2006). 10.1016/j.vaccine.2005.07.064

26 Kraan, H., van der Geest, A., Oosterhoff, D., Kruiswijk, C. & Soema, P. A Randomized, Double-Blind, Placebo-Controlled Phase I Study to Evaluate the Safety, Tolerability, and Immunogenicity of an Outer Membrane Vesicle (OMV) Platform-Based Vaccine Administered Intranasally to Healthy Adults. Vaccines (Basel) 14 (2026). 10.3390/vaccines14070575

27 Corman, V. M. et al. Detection of a novel human coronavirus by real-time reverse-transcription polymerase chain reaction. Euro Surveill 17 (2012). 10.2807/ese.17.39.20285-en

28 Wang, C. et al. Antigenic structure of the human coronavirus OC43 spike reveals exposed and occluded neutralizing epitopes. Nat Commun 13, 2921 (2022). 10.1038/s41467-022-30658-0

29 Wang, C. et al. A human monoclonal antibody blocking SARS-CoV-2 infection. Nat Commun 11, 2251 (2020). 10.1038/s41467-020-16256-y

30 Widjaja, I. et al. Towards a solution to MERS: protective human monoclonal antibodies targeting different domains and functions of the MERS-coronavirus spike glycoprotein. Emerg Microbes Infect 8, 516–530 (2019). 10.1080/22221751.2019.1597644

31 Cohen, A. A. et al. Mosaic RBD nanoparticles protect against challenge by diverse sarbecoviruses in animal models. Science 377, eabq0839 (2022). 10.1126/science.abq0839

32 Wec, A. Z. et al. Broad neutralization of SARS-related viruses by human monoclonal antibodies. Science 369, 731–736 (2020). 10.1126/science.abc7424

33 Greaney, A. J. et al. Comprehensive mapping of mutations in the SARS-CoV-2 receptor-binding domain that affect recognition by polyclonal human plasma antibodies. Cell Host Microbe 29, 463–476 e466 (2021). 10.1016/j.chom.2021.02.003

34 Pinto, D. et al. Broad betacoronavirus neutralization by a stem helix-specific human antibody. Science 373, 1109–1116 (2021). 10.1126/science.abj3321

35 Zhou, P. et al. Broadly neutralizing anti-S2 antibodies protect against all three human betacoronaviruses that cause deadly disease. Immunity 56, 669–686 e667 (2023). 10.1016/j.immuni.2023.02.005

36 Hou, Y. J. et al. SARS-CoV-2 Reverse Genetics Reveals a Variable Infection Gradient in the Respiratory Tract. Cell 182, 429–446 e414 (2020). 10.1016/j.cell.2020.05.042

37 V’Kovski, P., Kratzel, A., Steiner, S., Stalder, H. & Thiel, V. Coronavirus biology and replication: implications for SARS-CoV-2. Nat Rev Microbiol 19, 155–170 (2021). 10.1038/s41579-020-00468-6

38 Sterlin, D. et al. IgA dominates the early neutralizing antibody response to SARS-CoV-2. Sci Transl Med 13 (2021). 10.1126/scitranslmed.abd2223

39 Lapuente, D. et al. Protective mucosal immunity against SARS-CoV-2 after heterologous systemic prime-mucosal boost immunization. Nat Commun 12, 6871 (2021). 10.1038/s41467-021-27063-4

40 Winklmeier, S. et al. Intramuscular vaccination against SARS-CoV-2 transiently induces neutralizing IgG rather than IgA in the saliva. Front Immunol 15, 1330864 (2024). 10.3389/fimmu.2024.1330864

41 Adler, J. M. et al. An intranasal live-attenuated SARS-CoV-2 vaccine limits virus transmission. Nat Commun 15, 995 (2024). 10.1038/s41467-024-45348-2

42 Leekha, A. et al. Multi-antigen intranasal vaccine protects against challenge with sarbecoviruses and prevents transmission in hamsters. Nat Commun 15, 6193 (2024). 10.1038/s41467-024-50133-2

43 Darling, T. L. et al. Mucosal immunization with ChAd-SARS-CoV-2-S prevents sequential transmission of SARS-CoV-2 to unvaccinated hamsters. Sci Adv 10, eadp1290 (2024). 10.1126/sciadv.adp1290

44 Dowell, A. C. et al. Nasal mucosal IgA levels against SARS-CoV-2 and seasonal coronaviruses are low in children but boosted by reinfection. J Infect 87, 403–412 (2023). 10.1016/j.jinf.2023.08.013

45 Liu, Z. et al. Precision-engineered STING agonist nanoparticles enable coordinated mucosal-systemic immunity for durable pan-beta-coronavirus protection. Nat Nanotechnol 21, 1008–1021 (2026). 10.1038/s41565-026-02188-z

46 Thornton, V. et al. Safety and immunogenicity of New Zealand strain meningococcal serogroup B OMV vaccine in healthy adults: beginning of epidemic control. Vaccine 24, 1395–1400 (2006). 10.1016/j.vaccine.2005.09.043

47 Vesikari, T. et al. Immunogenicity and safety of an investigational multicomponent, recombinant, meningococcal serogroup B vaccine (4CMenB) administered concomitantly with routine infant and child vaccinations: results of two randomised trials. Lancet 381, 825–835 (2013). 10.1016/S0140-6736(12)61961-8

48 Chen, J. et al. Bat-infecting merbecovirus HKU5-CoV lineage 2 can use human ACE2 as a cell entry receptor. Cell 188, 1729–1742 e1716 (2025). 10.1016/j.cell.2025.01.042

49 Wang, N. et al. A MERS-CoV-like mink coronavirus uses ACE2 as an entry receptor. Nature 642, 739–746 (2025). 10.1038/s41586-025-09007-w

50 Xiong, Q. et al. ACE2-using merbecoviruses: Further evidence of convergent evolution of ACE2 recognition by NeoCoV and other MERS-CoV related viruses. Cell Insight 3, 100145 (2024). 10.1016/j.cellin.2023.100145

51 Allen, J. D. et al. Site-Specific Steric Control of SARS-CoV-2 Spike Glycosylation. Biochemistry 60, 2153–2169 (2021). 10.1021/acs.biochem.1c00279

52 Pino, P. et al. Trimeric SARS-CoV-2 Spike Proteins Produced from CHO Cells in Bioreactors Are High-Quality Antigens. Processes 8, 1539 (2020).

53 Zielinska, E. et al. Development of an improved microneutralization assay for respiratory syncytial virus by automated plaque counting using imaging analysis. Virol J 2, 84 (2005). 10.1186/1743-422X-2-84

54 Zettl, F. et al. Rapid Quantification of SARS-CoV-2-Neutralizing Antibodies Using Propagation-Defective Vesicular Stomatitis Virus Pseudotypes. Vaccines (Basel) 8 (2020). 10.3390/vaccines8030386

55 Ogoti, B. M. et al. Biphasic MERS-CoV Incidence in Nomadic Dromedaries with Putative Transmission to Humans, Kenya, 2022-2023. Emerg Infect Dis 30, 581–585 (2024). 10.3201/eid3003.231488

